# Nieman-Pick Type C2 proteins in *Aedes aegypti*: Their Structure-Function relationships and Expression in Uninfected *versus* Virus-infected Mosquitos

**DOI:** 10.1101/2021.11.15.468632

**Authors:** Prathigna Jaishankar Thambi, Cassandra M. Modahl, R. Manjunatha Kini

**Affiliations:** PES University, Bengaluru, Karnataka, India; Department of Biological Sciences, Faculty of Science, National University of Singapore, Singapore; Department of Pharmacology, Yong Loo Lin School of Medicine, National University of Singapore, Singapore; Department of Biochemistry and Molecular Biology, VCU School of Medicine, Virginia Commonwealth University, Richmond, Virginia, USA

**Keywords:** Dengue, Chikungunya, Zika, Differential expression, Chemoattractant, Promoter region, *cis* elements

## Abstract

*Aedes aegypti* is a major vector that transmits arboviruses through the saliva injected into the host. Salivary proteins help in uninterrupted blood intake and enhance the transmission of pathogens. We studied Nieman-Pick Type C2 (NPC2) proteins, a superfamily of saliva proteins that play important role in arbovirus infections. In vertebrates, a single conserved gene encodes for the NPC2 protein that functions in cholesterol trafficking. Arthropods, in contrast, have several genes that encode for divergent NPC2 proteins. We compared the sequences of 20 *A. aegypti* NPC2 proteins to the cholesterol-binding residues of human and bovine, and fatty acid-binding residues of ant NPC2 proteins. We identified four and one mosquito NPC2 proteins as potential sterol- and fatty acid-binding proteins, respectively. From the published data, we analysed the expression of NPC2 genes in various tissues and their differential expression in midgut and salivary gland post-arbovirus infections. NPC2 genes are downregulated rather than upregulated in virus-infected tissues. Interestingly, AAEL012064 is the only gene that is downregulated in both the midgut and salivary gland in all virus infections. In addition, AAEL001650 is downregulated in the salivary gland infected with CHIKV, DENV2 or ZIKV. This gene in the midgut is downregulated infected with DENV1 but upregulated with DENV2. We studied the variation in *cis* elements in the promoter regions of two groups of closely related NPC2 genes and the expression of relevant transcription factors (TFs). In the midgut infected with DENV1 or DENV2, six TFs (CRE-BP1, AP1, c-Jun, c-Fos, Odd, and NF-kB) appear to play an opposing role in the expression of AAEL006854. Two TFs (RXR-beta/alpha and USF) have a potential role in the downregulation of AAEL009556 in CHIKV-infected midgut and salivary gland. Similarly, two TFs (COUP and Ftz) may have a key role in the downregulation of AAEL009555 and AAEL009556 in DENV2-infected salivary gland.

## Introduction

Arbovirus infections spread vastly across the world and millions of individuals are affected every year. Mosquitoes act as vectors in transmitting several viruses, including Dengue (DENV), Chikungunya (CHIKV), Zika (ZIKV), Yellow fever, West Nile, and Japanese encephalitis leading to deadly diseases. These diseases induce a wide range of clinical symptoms ranging from mild fever and headache to more severe seizures and fatal haemorrhage [1]. DENV is endemic and epidemic in the tropical and subtropical regions and causes around 390 million infections per year [1]. CHIKV first appeared in Africa in the 1950s, re-emerged in 2004 and accounted for the largest outbreak, infecting over 6 million people [2]. The sudden onset of high-grade fever, polyarthralgia, rash, and a headache characterize this illness [2]. ZIKV, the most recent arbovirus in the news, has led to over 1.5 million infections in the past five years [3]. It is the first arbovirus that can be transmitted sexually in humans [4]. ZIKV infections are associated with Guillain-Barre syndrome affecting 2 in 100,000 people and microcephaly affecting 1 in 10 babies born to mothers who contracted the infection during pregnancy [5]. The frequent re-emergence of arbovirus infections and the long-lasting effect of symptoms have made them a global health concern.

*Aedes aegypti* and *A. albopictus* are primary or secondary vectors of DENV, CHIKV and ZIKV [4]. In a gonotrophic cycle, female mosquitoes feed on host blood to obtain the nutrients necessary for producing viable eggs. During these blood meals, a ‘naïve’ mosquito acquires the pathogens circulating in the infected host blood. These pathogens initially reach the midgut, get past its physical and immune barriers and multiply numerous times [6]. The presence of viral proteins and RNA can be seen in the midgut as early as 2-3 days after an infected blood meal, although it peaks around ten days [7]. The virus then spreads to other regions of the body, eventually invading the salivary gland. Subsequently, the pathogens in the saliva are transferred to a healthy person during the next blood meal infecting him/her.

During a bite, mosquito injects its saliva into the host to overcome host responses to vessel injury such as blood coagulation, platelet aggregation, and vasoconstriction [8]. The infusion of saliva ensures a continuous blood flow during the meal. Inadvertently, the saliva infusion aids in the transfer of viruses. Similar to saliva proteins from other hematophagous arthropods, mosquito saliva proteins enhance the infectivity of the pathogens [8]. Thus, mosquito saliva proteins facilitate both blood intake and enhanced pathogen transmission. As a first step to understand their role, the saliva proteins have been profiled using transcriptomes and proteomes [9–14]. The saliva proteins are classified into several superfamilies based on their similarity to various characterized proteins [15]. Each protein superfamily contains multiple isoforms encoded by distinct genes with varied sequences. Although these proteins share comparable disulphide pairings and three-dimensional structures, they may exhibit distinct functions. Only a small number of mosquito proteins have been characterized for their structure, function, and mechanism of action so far. The physiological functions of most of other proteins are yet to be established.

Members of one superfamily of mosquito saliva proteins share sequence similarity with the human Niemann-Pick Type C2 (NPC2) protein, and hence classified as NPC2 protein family. Human NPC2 is a small protein found in the lysosome, belonging to the MD-2 lipid recognition domain family [16]. NPC2 and NPC1 (structurally unrelated to NPC2) are involved in binding, trafficking, and distribution of free cholesterol out of lysosomes to other parts of the cell and contribute to cholesterol homeostasis in all cell types [16]. Mutations in either NPC1 or NPC2 protein can cause accumulation of cholesterol in lysosomal compartments leading to a rare neurovisceral lipid storage disorder, Nieman Pick Disease Type C [16]. This rare autosomal recessive disorder affects 1 in 100,000 people and can be fatal [17].

A single highly conserved gene encodes the NPC2 protein in mammals and other vertebrates, with identity of 75% among mammals and 55-70% across vertebrates [18]. The high conservation of NPC2 proteins is probably associated with its single functionality. Unlike vertebrates, arthropods have several genes encoding structurally divergent NPC2 proteins. In *A. aegypti*, 13 NPC2 proteins are expressed in the salivary gland [12]. However, a genome-wide search indicates that there are 20 NPC2 genes (Genome version AaegL5.3). These NPC2 proteins probably play distinct biological roles. The structure-function relationships and the physiological roles of the NPC2 proteins are poorly understood. In this paper, we have compared *A. aegypti* NPC2 proteins to human and bovine NPC2 proteins that bind to cholesterol and ant NPC2 that binds to fatty acid. Based on the similarities, we have identified NPC2 proteins that may bind sterols and fatty acid, respectively.

In addition to salivary gland, NPC2 genes are expressed in the midgut, brain, proboscis, and antenna of mosquitos [9–14]. The transcriptomes of *A. aegypti* midgut from uninfected mosquitoes with those infected with CHIKV, DENV1 or DENV2 were compared in three different studies [9–11]. In a recent study, we compared the transcriptomes of *A. aegypti* salivary gland from uninfected mosquitos with those infected with CHIKV, DENV2, or ZIKV [12]. Although the expression of NPC2 genes is affected by the infection of various arboviruses, these data were not analysed for differential expression of NPC2 genes. Thus, little is known of their role in arbovirus infections. Jupatanakul et al. suggest an agonist role for these lipid-binding proteins in DENV2 infection [19]. So far, no other studies have evaluated the roles of NPC2 genes in mosquitos infected with DENV1, CHIKV, or ZIKV. Here, we have focused on the analyses of the differential expression of NPC2 genes in the midgut and salivary gland of *A. aegypti* infected with DENV1/DENV2, CHIKV, or ZIKV. By studying the transcription factor (TF) binding sites in the proximal promoter region (PPR) and the expression of TFs in relevant tissues, we have identified TFs that may affect the expression of two sets of closely related NPC2 genes.

## Materials and methods

### Analyses of nucleotide and protein sequences of NPC2 genes

The DNA, mRNA and protein sequences for 20 *A. aegypti* NPC2 genes were downloaded from VectorBase [20]. The protein sequences were analysed for the signal peptides using SignalP 5.0 [21]. The nucleotide and protein sequences were aligned using the Clustal Omega bioinformatics tool [22]. The similarities between mRNAs and proteins were visualised as heatmaps using Matplotlib [23]. Genome wide distribution of NPC2 genes on the three chromosomes of *A. aegypti* was analysed to understand the location of genes and identify the gene clusters in each chromosome. The relationship between the distance among genes within the cluster and their sequence similarities were analysed. Intron-exon profiles were studied to identify the presence of retrogenes. We also examined synteny between two genes sharing the highest identity.

The evolutionary history was inferred by using the Maximum Likelihood method and Whelan And Goldman model [24]. The bootstrap consensus tree inferred from 1000 replicates [25] is taken to represent the evolutionary history of the taxa analyzed [25]. Branches corresponding to partitions reproduced in less than 50% bootstrap replicates are collapsed. The percentage of replicate trees in which the associated taxa clustered together in the bootstrap test (1000 replicates) are shown next to the branches [25]. Initial tree(s) for the heuristic search were obtained automatically by applying Neighbor-Join and BioNJ algorithms to a matrix of pairwise distances estimated using the JTT model, and then selecting the topology with superior log likelihood value. A discrete Gamma distribution was used to model evolutionary rate differences among sites (5 categories (+*G*, parameter = 7.8010)). The rate variation model allowed for some sites to be evolutionarily invariable ([+*I*], 3.86% sites). This analysis involved 23 amino acid sequences, and were first aligned using MUSCLE. There was a total of 179 positions in the final dataset. The human 5KWY was used to root the tree. All analyses were conducted in MEGA X [26], [27].

### Structure-function relationships of NPC2 proteins

To identify the residues in *A. aegypti* NPC2 proteins involved in the interaction with cholesterol or fatty acid, we compared their sequences with the amino acid residues that are involved in binding to cholesterol or fatty acid in the crystal structures of reference proteins, human (PDB 5KWY), bovine (PDB 2HKA), and ant NPC2 (PDB 3WEB) complexed with cholesterol or fatty acid [28–30]. The structures were visualised and analysed using PyMOL 2.5. The residues of our proteins were compared to the binding residues of reference proteins using heatmaps created with Matplotlib [23]. The identical residues were marked in green, conserved residues in yellow and non-conserved residues in red. This provided a simple visual scheme to identify NPC2 proteins that would potentially bind sterol or fatty acid.

### Expression of NPC2 genes in mosquito tissues

The expression profiles of NPC2 genes were evaluated using reads from transcriptomic studies [9–14]. The raw reads of uninfected female *A. aegypti* are available for the midgut, salivary gland, antenna, brain, proboscis and whole body [9–14]. The mosquitos were blood-fed in the transcriptome studies of the midgut, salivary gland and whole body, whereas non-blood fed mosquitos were used for the transcripts from antenna, brain and proboscis. We aligned and quantified the raw reads of RNAseq data from the midgut, antenna, brain, proboscis, and whole-body using Bowtie2 and RSEM to evaluate the expression of NPC2 genes. For the salivary gland, the reads were normalised using DESeq2 [12]. Heatmaps to represent the rank of expression of highly expressed genes in each tissue were created using Matplotlib [23].

### Differential expression of NPC2 genes in infected mosquito

The raw reads of transcripts from only the midgut and salivary gland of infected female *A. aegypti* are available [9–12]. The mosquitos were infected orally with CHIKV, DENV1 or DENV2 in the midgut and dissected at 1-day post infection (dpi), 4-dpi and 14-dpi for CHIKV, DENV1, or DENV2 respectively [9–11]. In contrast, salivary gland was infected orally with CHIKV, DENV2 or ZIKV and dissected at 7-dpi for CHIKV and 14-dpi for DENV2 or ZIKV [12]. Genes that showed a differential expression of over 20% of the control in the midgut and salivary gland were analysed. A simple representation of the differential expression allowed us to effectively compare the expression pattern between genes and tissues.

### Transcription factors that regulate expression of NPC2 genes

The PPR sequences (600 bp upstream from the transcription initiation site) for two groups of closely related genes – Group A (AAEL006854 and AAEL020314) and Group B (AAEL009553, AAEL009555, and AAEL009556) were collected from VectorBase [20]. The TF binding sites (*cis* elements) in these sequences were predicted using TRANSFAC 4.0 and AliBaba 2 [31], [32]. We analysed the reads for the expression of relevant TFs in the midgut and salivary gland of uninfected and infected mosquitos obtained from the previous transcriptomic studies [9–12]. TFs that showed a differential expression of over 20% of the control in the midgut and salivary gland were analysed. Heatmaps for differential expression of the TFs from this analysis was plotted using Matplotlib to identify the TFs that exhibit a high degree of differential expression.

## Results

### Sequence analysis

To identify whether the *A. aegypti* NPC2 proteins are secreted or cytosolic proteins, we analysed them for the presence of signal peptide. All *A. aegypti* NPC2 proteins have signal peptides at the N-terminal, and hence, they are secreted proteins. We aligned mature amino acid sequences of *A. aegypti* NPC2 proteins with *Homo sapiens* (NP_006423.1), *Bos taurus* (NP_776343.1), and *Camponotus japonicus* (BAO48214.1) NPC2 proteins (Figure 1A). The mature *A. aegypti* NPC2 protein sequences ranged in length from 130 to 146 amino acid residues (Figure 1A). AAEL026174 is the shortest, while AAEL025109 is the longest. Except for AAEL004120, which has eight cysteines, all the proteins have six conserved cysteines that form three disulphide linkages C1-C6, C2-C3, C4-C5 (Figure 1A). Eight NPC2 proteins that have at least one *N*-glycosylation site are AAEL001654, AAEL0026174, AAEL015137, AAEL012064, AAEL006854, AAEL020314, AAEL004120, and AAEL025109 (Figure 1A). Human and bovine NPC2 have one *N*-glycosylation site each, whereas ant NPC2 and remaining 12 *A. aegypti* NPC2 proteins do not have any *N*-glycosylation sites (Figure 1A).

**Fig 1.**
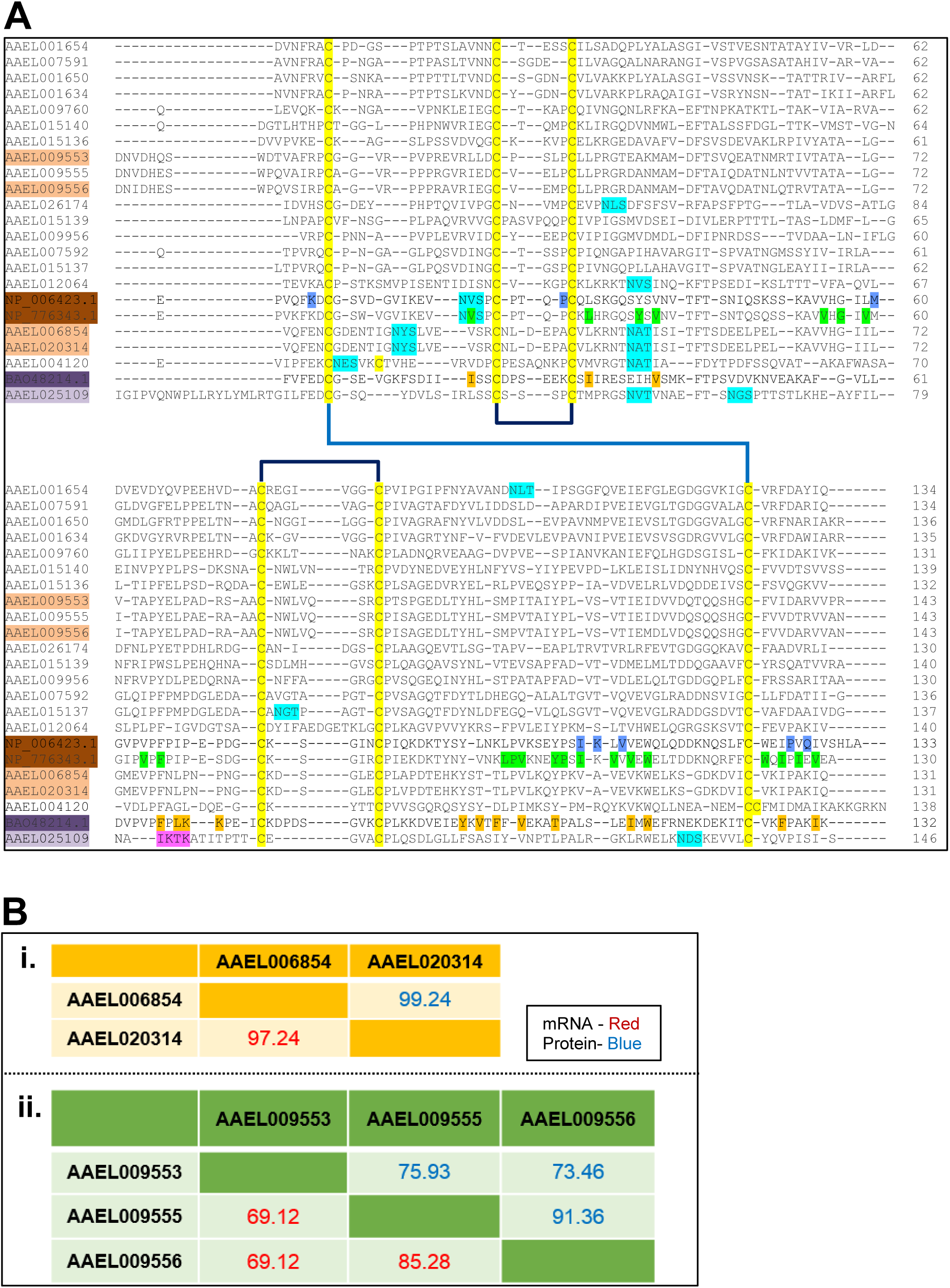
Sequence alignment of mature *Aedes aegypti* NPC2 proteins. **(A)** The mature NPC2 proteins of *Aedes aegypti* (AAEL0xxxxx)*, Homo sapiens* (NP_006423.1), *Bos taurus* (NP_776343.1) and *Camponotus japonicus* (BAO48214.1) were aligned using Clustal Omega. Six conserved cysteines are highlighted in yellow and the three predicted disulphide pairings (C1-C6, C2-C3, C4-C5) are shown. The predicted *N*-Glycosylation sites are highlighted in cyan. The residues involved in cholesterol binding in bovine NPC2 are highlighted in green, while the residues interacting with NPC1 and hence, cholesterol transfer, in human NPC2 are highlighted in blue. The accession numbers of cholesterol-binding bovine and human NPC2 are highlighted in dark brown, while those of predicted *A. aegypti* NPC2 proteins are highlighted in light brown. The residues involved in fatty acid binding in ant NPC2 are highlighted in orange. The accession numbers of fatty acid-binding ant NPC2 and predicted *A. aegypti* NPC2 are highlighted in dark lavender and light lavender, respectively. The sequence pattern IKTK that functions as the double lysine site in AAEL025109 is highlighted in purple. **(B)** The mRNA and protein identity of closely related genes belonging to groups A (i) and B (ii) NPC2 genes are shown.

The sequence identity of mature proteins ranged from 14-99% (Figure S1). The proteins with the highest identity AAEL006854 and AAEL020314 are 131 amino acid residues in length and differ only by one residue; alanine89 in AAEL006854 replaced by valine89 in AAEL020314 (Figure 1B(i)). AAEL009555 and AAEL009556 also show high identity of 91.36% (Figure 1B(ii)). The gene identity for this pair is 75.79%. AAEL025109 is the most distinct NPC2 protein, with only 14-25% identity to the others (Figure S1). AAEL007591 and AAEL025109 are the least identical proteins with 14% identity. Although human and bovine NPC2 share 80% identity, they share only 18-40% identity with *A. aegypti* NPC2 proteins, while the ant protein shares 14-37% identity with *A. aegypti* NPC2 proteins.

### Genome wide distribution

NPC2 genes are distributed among the three chromosomes of *A. aegypti*, except for AAEL020314 gene, which is located on an extra-chromosomal scaffold (Figure 2A). Ten genes are located on chromosome 1. Interestingly, nine of them form a cluster and lie within 1.6 Mbp - AAEL007592, AAEL007591, AAEL015137, AAEL015139, AAEL009956, AAEL0026174, AAEL015136, AAEL015140, and AAEL025109. Remarkably, AAEL009956 and AAEL026174 are the closest with an intergenic distance of only 40 bp (Figure 2A). Followed by AAEL015139 and AAEL009956 (separated by 88 bp), and AAEL015136 and AAEL015140 (separated by 288 bp). Interestingly, despite their proximity these genes share only low identity between their mRNA (35-53%) and protein (13-47%) sequences with the exception of AAEL007592 and AAEL015137, which share over 73% identity (Figure S1). The last gene AAEL025109 is unusually close to the telomere with only 7.7 Mbp from the end of the chromosome. Four genes are found on chromosome 2, where three genes are clustered together AAEL001654, AAEL001650, and AAEL001634 (Figure 2A). Even though these genes are in a cluster, they only share about 58-72% mRNA identity and 49-69% protein identity (data not shown). Five genes are found on chromosome 3, including one cluster of three genes AAEL009553, AAEL009555, and AAEL009556 (Figure 2A). Two genes AAEL009555 and AAEL009556 are 280 kbp apart and have 85.28% and 91.36% mRNA and protein identity, respectively (Figure 1B(ii)). AAEL009553 shares 69.12% and 69.12% mRNA and 75.93% and 73.46% protein identity with AAEL009555 and AAEL009556, respectively (Figure 1B(ii)).

**Fig 2.**
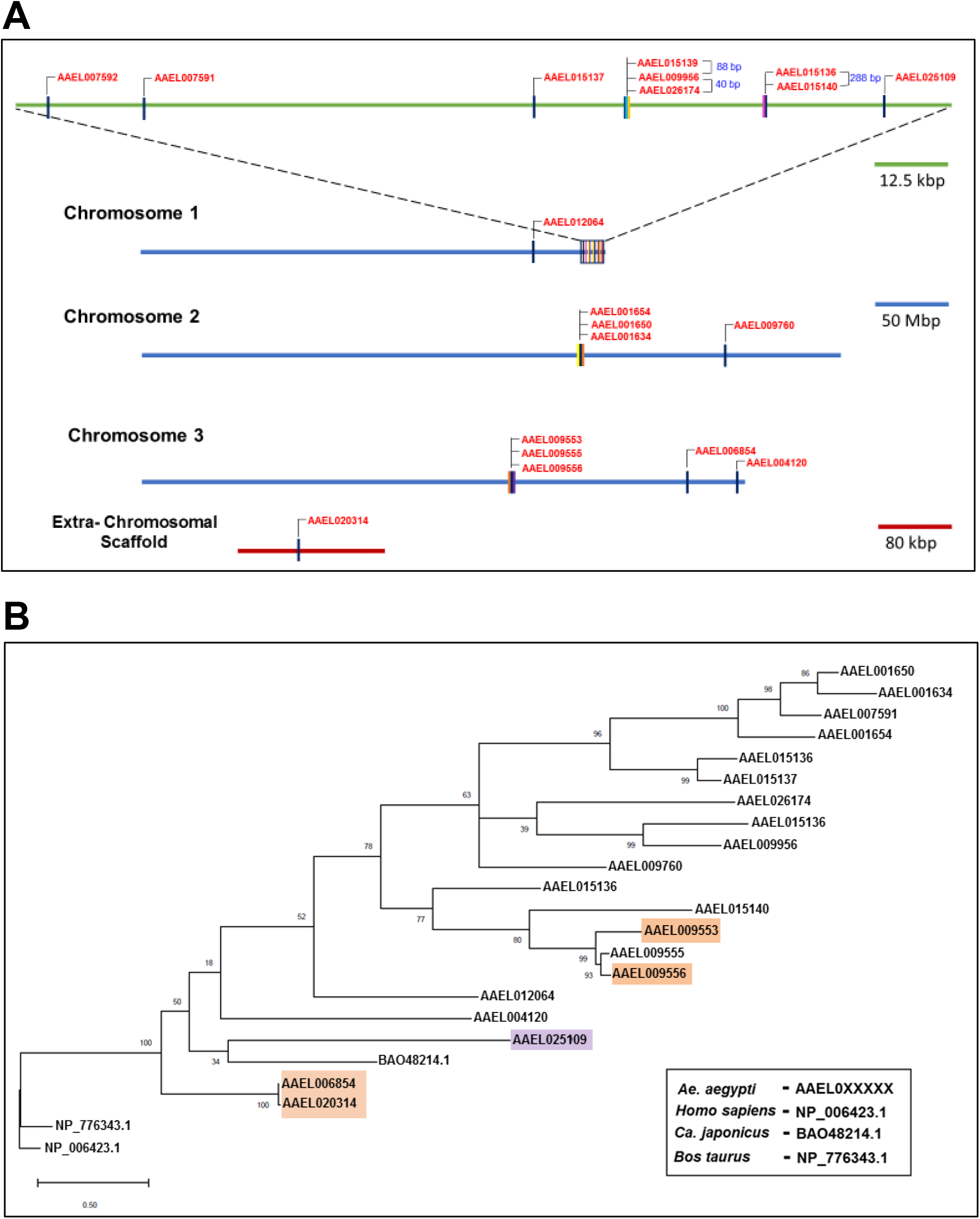
Genomic distribution and phylogenetic relationships of *A. aegypti* NPC2 genes. **(A)** Nineteen *A. aegypti* NPC2 genes are distributed among the three chromosomes, while one gene is located on an extra-chromosomal scaffold. The genome distribution is made to scale. Ten genes are located on chromosome 1, with nine genes forming a cluster. Within this cluster, two groups of three and two genes are separated by unusually small intergenic segments (indicated in bp). Four genes are located on chromosome 2, with three genes forming a cluster. Five genes are distributed on chromosome 3, with three genes forming a cluster. **(B)** Phylogenetic tree of *A. aegypti* NPC2 proteins along with Bovine (2HKA), Human (5KWY), and Ant (3WEB) NPC2 proteins were constructed using SeaView, PhyML and Figtree. This tree shows the diversity of the mosquito NPC2. The four potential cholesterol-binding proteins are highlighted in brown, while the one potential fatty acid-binding protein is highlighted in lavender.

### Intron-exon profiles of NPC2 genes

We analysed intron-exon profiles, chromosome location, gene and protein sizes, and the direction of transcription of all NPC2 genes (Figure S3). Most genes are less than 1200 bp long, ranging from 465 to 1135 bp. AAEL012064, AAEL020314, AAEL006854, AAEL025109, and AAEL009760 genes, ranging from 14,000 to 41,500 bp, are the exceptions (Figure S3). The shortest and the longest genes with lengths 465 bp and 41,500 bp are AAEL015137 and AAEL012064, respectively. The intron-exon profiles are also varied. Fourteen genes contain only one intron and two exons (Figure S3). Only one gene (AAEL004120) has two introns and three exons, and three genes (AAEL025109, AAEL006854 and AAEL020314) contain three introns and four exons (Figure S3). AAEL012064 has the highest number of introns (five), two of them are in the 5’ UTR. Only AAEL015137 has no introns. Since it does not contain a poly (A) tail, it is most likely not a retrogene. Interestingly, AAEL020314 (extrachromosomal) and AAEL006854 that share high protein identity (99.24%), but do not share the synteny. There was no matching identity in the vicinity of these two genes (data not shown).

Phylogenetic analysis was conducted on mature protein sequences (Figure 2B). The potential cholesterol-binding proteins are highlighted in rust and the potential fatty-acid binding protein in lavender (Figure 2B). Potential cholesterol-binding AAEL006854 and AAEL020314 are evolutionarily distinct from all other *A. aegypti* NPC2 proteins, including the other two putative cholesterol-binding AAEL009553 and AAEL009556. Although still closely related to other *A. aegypti* NPC2 proteins, the putative fatty acid-binding AAEL025109 clustered with the ant fatty acid-binding NPC2 (Figure 2B).

### Structure - Function relationships of NPC2 proteins

#### NPC2 proteins in cholesterol binding and transport

As mentioned in the Introduction, the mammalian NPC2 proteins play a pivotal role in cholesterol binding and egress of the lipid from the lysosome to other organelles [16]. Bovine NPC2 (PDB 2HKA) protein has an immunoglobulin-like fold stabilised by three disulphide bridges. Apo NPC2 (PDB 1NEP) has multiple small loosely packed hydrophobic cavities which are closed. When an appropriate ligand, such as cholesterol, is bound, a deep hydrophobic cavity flanked by two β strands expand by repositioning the sidechains of several residues to fit the ligand [30]. This malleability is a characteristic of vertebrate NPC2 proteins, unlike many lipid-binding proteins, which have large pre-existing cavities to bind various ligands [30].

The cholesterol-3-*O*-sulphate molecule bound to a predominantly hydrophobic sterol binding tunnel of bovine NPC2 is formed by 22 amino acid residues, V20, L30, Y36, V38, V55, G57, V59, V64, F66, L94, P95, V96, Y100, P101, I103, V105, V107, W109, W122, I124, I126, and V128 [30] (Figure 3B). The side chains of ten residues reposition when cholesterol is bound, L30, Y36, V38, F66, L94, P95, V96, Y100, P101, and I103 [30]. The entrance of this tunnel is formed by six residues – V59, V64, F66, Y100, P101 and I103 [30]. Cholesterol sulphate interacts with all these residues. Particularly, the two aromatic F66 and Y100 residues are involved in most interactions. Mutation of these aromatic residues to aliphatic Alanine residue severely compromises the cholesterol binding activity [33]. Thus, the aromaticity of these residues contributes to protein structural stability as well as to the strong hydrophobic interactions with cholesterol. Similarly, replacement of V96, located in the middle of the cavity, with Phenylalanine hinders the cholesterol binding due to steric clash [33]. On the other hand, V64F and W122A mutations has little impact on ligand binding [33]. P101 and V20, situated at the top and the base of the hydrophobic tunnel (Figure 3B), are crucial residues that form hydrophobic interactions with cholesterol. In Nieman-Pick disorder patients, P101S and V20M mutations lead to loss of function [34].

**Fig 3.**
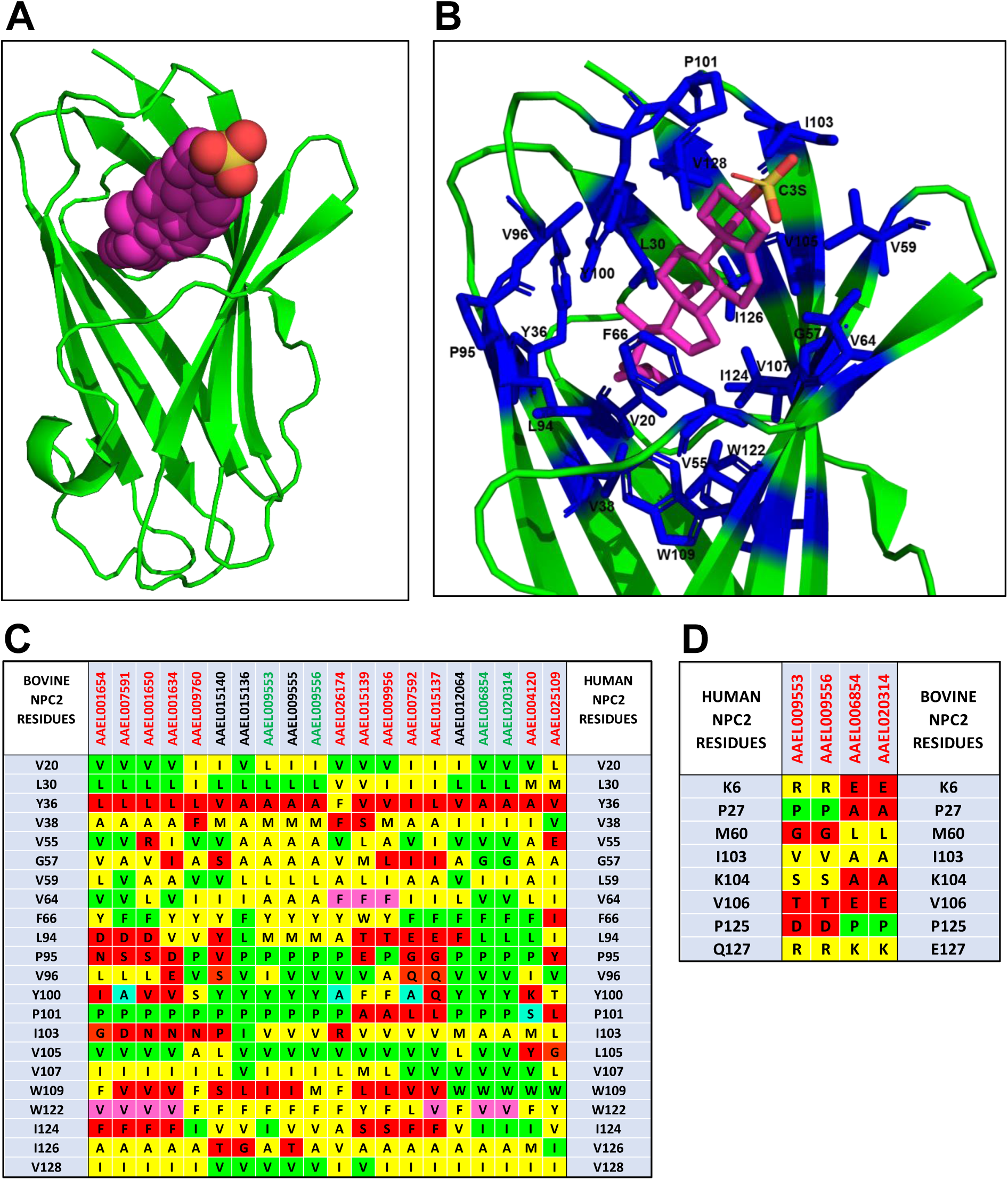
Cholesterol binding site in NPC2 proteins. **(A)** Ribbon representation of bovine NPC2 bound to cholesterol sulphate (2HKA) are shown using PyMOL. Cholesterol sulphate is depicted as a stick model. Carbon, magenta; oxygen, red; and sulphur, yellow. The residues involved in forming the predominantly hydrophobic tunnel are depicted as stick model (blue). **(B)** Comparison of amino acid residues of *A. aegypti* NPC2 proteins with those in cholesterol binding site of bovine NPC2. Identical residues are highlighted in green, conserved residues in yellow, and non-conserved residues in red. Mutations studies indicated that the residues highlighted in teal show severe reduction, while those in violet have little to no impact on cholesterol binding. **(C)** Comparison of amino acid residues of four *A. aegypti* potential cholesterol-binding NPC2 proteins with those in human NPC2 involved in interaction with NPC1. Identical residues are highlighted in green, conserved residues in yellow, and non-conserved residues in red.

All mosquito NPC2 proteins were analysed for the 22 amino acid residues involved in cholesterol binding. In Figure 3C, we have highlighted all identical residues in green, conserved residues in yellow, and non-conserved residues in red. In all *A. aegypti* NPC2 proteins, aliphatic residues V20, L30, V59, and V107 are either conserved or replaced with other aliphatic amino acid residues (Figure 3C). The Y100A mutation, which results in loss of cholesterol binding [33], is found in AAEL007591, AAEL007592, and AAEL026174 (Figure 3C). Similarly, the P101S mutation is found in AAEL004120 and leads to loss of cholesterol binding [34]. Thus, these four proteins probably do not bind cholesterol or related molecules. The mutations V20M, F66A, and V96F that also results in a loss of cholesterol binding [34] are not found in *A. aegypti* NPC2 proteins. We speculate that the F66I mutation found in AAEL025109 (Figure 3C), similar to F66A, may lead to a loss of cholesterol binding. Twelve proteins, AAEL001654, AAEL007591, AAEL001650, AAEL001634, AAEL009760, AAEL015139, AAEL009956, AAEL007592, AAEL015137, AAEL015140, AAEL009555 and AAEL025109 have one or more polar substitutions in the hydrophobic tunnel making it unsuitable for cholesterol binding (Figure 3C). Further, the replacement of P95 or P101 in these proteins can substantially compromise their conformation. Hence, these twelve proteins are probably not involved in cholesterol binding. On the other hand, the mutation V64F that had little impact on cholesterol binding [33] was found in AAEL026174, AAEL015139, and AAEL009956 (Figure 3C). In AAEL009556, the cholesterol binding pocket is mostly intact except for a single mutation, Y36A (Figure 3C). AAEL006854 and AAEL020314, on the other hand, have two mutations: Y36A and W122V. As W122A showed to have little impact on cholesterol binding [33], we assume that the W122V mutation may also have little impact due to the aliphatic, hydrophobic nature of Alanine and Valine. AAEL009553 contains two mutations Y36A and W109I (Figure 3C). We speculate that the mutation of W109 to aliphatic, hydrophobic residue in the hydrophobic tunnel may have a limited effect on cholesterol binding. Based on the above analysis, we classify four proteins, AAEL009556, AAEL006854, AAEL020314, and AAEL009553 as potential cholesterol binding proteins.

After binding cholesterol, NPC2 interacts with NPC1 to form a cholesterol transfer complex [30]. Following this transfer, NPC1 transports cholesterol to other cell organelles to maintain the cholesterol homeostasis [30]. The crystal structure of Human NPC1-NPC2 complex (PDB 5KWY) reveals the residues involved in the interaction between NPC2 and the middle, luminally oriented domain (MLD) of NPC1. NPC1-MLD binds at the top of cholesterol-binding tunnel of NPC2 primarily through its two projecting loops [28]. The binding interface consists of residues that are involved in both hydrophobic and hydrophilic interactions. Eight NPC2 residues, K6, P27, M60, I103, K104, V106, P125, and Q127, form the binding site [28]. Four of these eight residues, K6, M60, K104, and Q127, have distinct orientations in apo and cholesterol bound structures. Interestingly, M60 is involved in hydrophobic and hydrophilic interaction with NPC1 [28]. We analysed all four potential cholesterol-binding proteins from *A. aegypti* for these eight residues. Seven-out-of-eight (87.5%) residues are mutated in each NPC2 protein; 50% of these mutations are conserved, while other 50% mutations are non-conserved (Figure 3D). The key residue M60 is replaced by residues Gly/Leu in all the proteins (Figure 3D). Aliphatic V106 is replaced by a polar (charged Glu or uncharged Thr) residue resulting in decreased hydrophobicity. P27A mutation is found in AAEL006854 and AAEL020314, whereas P125D mutation is found in AAEL009553 and AAEL009556 (Figure 3D). As unfavourable mutations are observed in all of the potential cholesterol-binding NPC2 proteins, we do not expect *A. aegypti* NPC2 proteins to interact with human NPC1-like proteins. Instead, they may interact with *A. aegypti* NPC1 proteins (AAEL009531 and AAEL019883). Both these proteins have non-conserved mutations (Figure S4) replacing all the six residues of human NPC1 (Q421, Y423, P424, F503, F504, Y506) that interact with NPC2 [28]. Due to such drastic changes in the interaction sites in both *A. aegypti* NPC1 and NPC2 proteins, we cannot speculate whether any of *A. aegypti* NPC2 proteins would form complex with *A. aegypti* NPC1 proteins and are involved in cholesterol transfer or not. It may be possible that *A. aegypti* NPC2 proteins bind sterols to perform other physiological functions. Interestingly, the pair NPC1 (AAEL009531) and NPC2 (AAEL001650) is suggested to function together and negatively affect the immune deficiency (IMD) immune signalling pathway in mosquitos infected with DENV2 [19].

#### NPC2 proteins in semiochemical binding and communication

NPC2 in arthropods help in recognising and binding of semiochemicals, thereby regulating the chemosensory activity [29], [35]. The NPC2 protein of ant *Camponotus japonicus* is expressed in the antenna and binds to various semiochemicals, such as long chain fatty acids and alcohols [29]. Similar to the mammalian NPC2, ant NPC2 has a β-sandwich fold. However, it has a large ligand-binding cavity in the interior [29].The crystal structure of ant NPC2-oleic acid complex (PDB 3WEB) reveals that the ligand is bent into a U-shaped conformation to fit into the binding pocket (Figure 4A). Interestingly, the polar carboxylic acid group of oleic acid is facing the bottom of the hydrophobic cavity, while the polar sulphate group of cholesterol sulphate is fully exposed at the top of the hydrophobic cavity (Figures 3A and 4A). The binding pocket of ant NPC2 is formed by 16 amino acid residues, namely I18, I30, V38, F66, L68, K69, K70, Y93, V95, F97, V99, T103, I110, W112, F127, and I131 (Figure 4B). To better accommodate the ligand, W112 has distinct orientations in the apo and bound structures i.e., it flips its side chain by 120º to avoid a steric clash with oleic acid. Ant NPC2 has a distinct double lysine site, K69 and K70 that forms two hydrogen bonds with O1 and O2 atoms of oleic acid [29]. These bonds are crucial for the high affinity binding of oleic acid [29]. K69P and K70N/Q mutations are commonly found in other arthropods and vertebrates. This significant change explains the low affinity of fatty acids to mammalian NPC2 [29].

**Fig 4.**
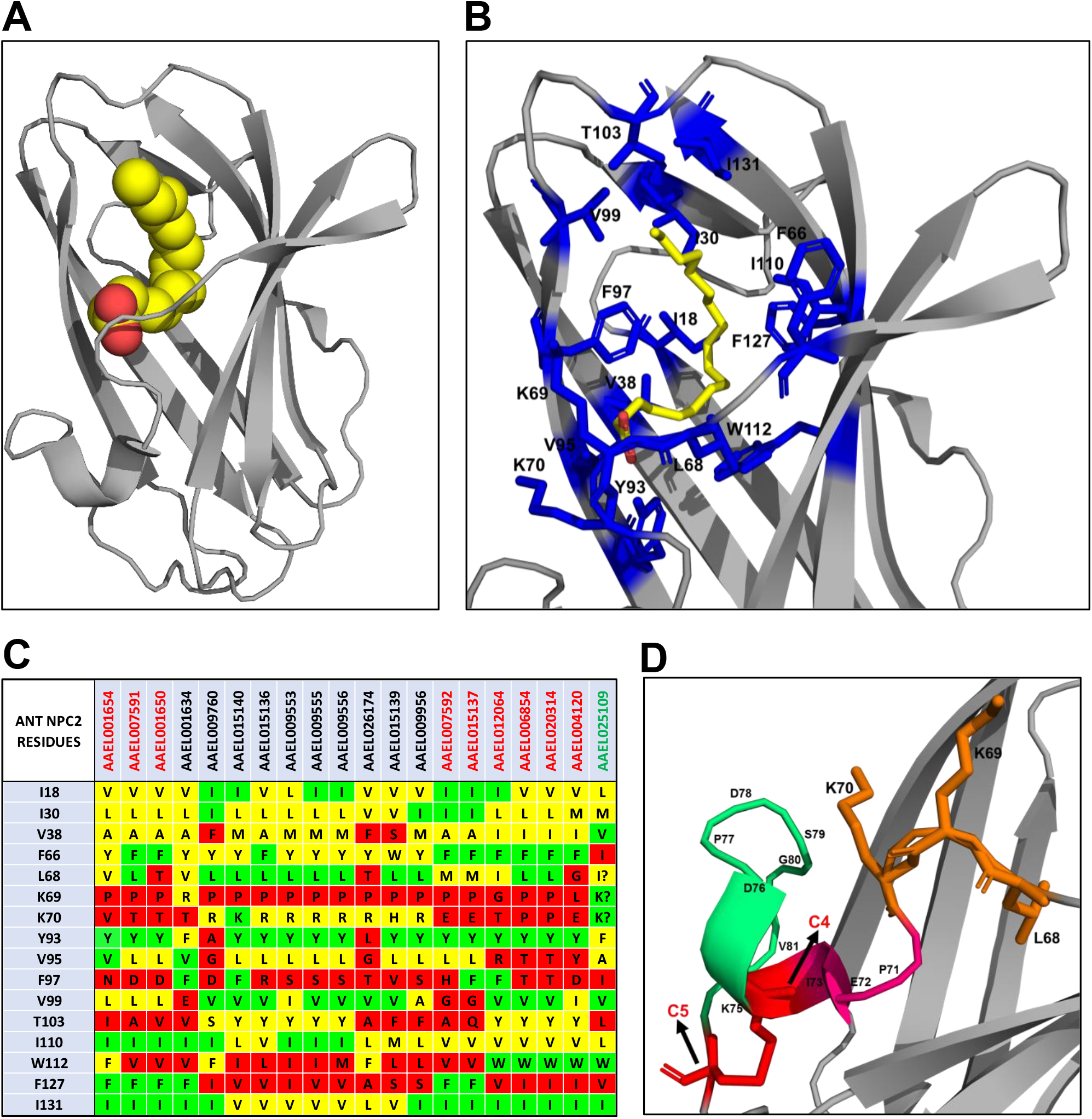
Fatty acid binding site in NPC2 proteins. **(A)** Ribbon representation of ant NPC2 bound to oleic acid (3WEB) are shown using PyMOL. Oleic acid is depicted as a stick model. Carbon, yellow and oxygen, red. The residues involved in forming the cavity are depicted as stick model (blue). **(B)** Comparison of amino acid residues of *A. aegypti* NPC2 proteins with those in fatty acid binding site of ant NPC2. Identical residues are highlighted in green, conserved residues in yellow, and non-conserved residues in red. **(C)** The segment of double lysine site that are in a loop in the Ant protein. In AAEL025109, the absence of series of prolines allows the beta strand to extend. The two lysine residues would then be on the same surface of the beta strand available to bind oleic acid.

All mosquito NPC2 proteins were analysed for the 16 amino acid residues involved in fatty acid binding. In all the proteins, three residues - I18, I30, and I131 are either conserved or replaced by conserved aliphatic amino acid residues (Figure 4C). W112, a residue that changes its orientation as mentioned above, is conserved in AAEL012064, AAEL006854, AAEL020314, AAEL004120, and AAEL025109 (Figure 4C), while it is replaced by F112 in AAEL001654, AAEL009760, and AAEL026174. In rest of the proteins, W112 is replaced by an aliphatic amino acid residue (Figure 4C). The double lysine site that is critical for oleic acid binding is mutated in most *A. aegpyti* NPC2 proteins. K69P mutant is found in 16 proteins, making it unfavourable for hydrogen bonding (Figure 4C). AAEL012064 and AAEL004120 have K69G and K69L mutations, respectively. K69R mutation is found in AAEL001634 (Figure 4C). On the other hand, K70 is mutated in 18 proteins. Seven proteins have a conserved residue mutation K70R and one protein has a mutation K70H (Figure 4C). The rest of the proteins have non-conserved mutations. K70 residue is conserved in AAEL015140 and AAEL025109. We speculate that the mutation of K69 or K70 to a polar, basic residue (Arg/ His) may allow the protein to form a single hydrogen bond with oleic acid, although the binding affinity could be affected. F97, that forms three hydrophobic interactions with the ligand, is replaced with a polar charged/uncharged or aliphatic amino acid residue in most of the proteins, resulting in reduction of hydrophobicity. Hence, we speculate that 19 of 20 mosquito NPC2 proteins may not be involved in fatty-acid binding. One protein, AAEL025109, contains four mutations F66I, F97I, T103L and F127V (Figure 4C). The two conserved K69 and K70 residues that are in a loop of ant NPC2 (Figure 4D) is replaced IKTK in a β-strand. The absence of series of proline residues (PVPFPLKKPE) in the vicinity allows the 6^th^ β strand to extend, subsequently allowing these two lysine residues to be available on the same surface of the β strand for hydrogen bonding with oleic acid. Based on this analysis, we conclude that AAEL025109 may be a potential fatty-acid binding protein.

### Expression of NPC2 genes in *A. aegypti* tissues

Analysis of the previously published transcriptomic data suggests that NPC2 genes are expressed in tissues like midgut, salivary gland, brain, antenna, and proboscis [9–14]. In uninfected female mosquitos, seven genes are significantly expressed in the midgut (Table S1A). The highly expressed genes in the midgut follow the order AAEL015136>> AAEL07591≥ AAEL009760>> AAEL026174> AAEL009556> AAEL007592>> AAEL015137 (Table S2A). Similarly, seven NPC2 genes are significantly expressed in salivary gland (Table S1B). The highly expressed genes in the salivary gland follow the order AAEL004120>> AAEL012064>>> AAEL006854>> AAEL009760>> AAEL001634> AAEL001650 = AAEL015136 (Table S2B). AAEL015136 and AAEL004120 shows the highest expression in the midgut and salivary gland, respectively (Table S2A and S2B). Interestingly, AAEL015136 that shows the highest expression in the midgut, is expressed in low amounts in the salivary gland.

Ten genes (AAEL001634, AAEL007591, AAEL009555, AAEL009956, AAEL015136, AAEL015137, AAEL015139, AAEL015140, AAEL026174 and AAEL025109) are not expressed in the antenna, brain and proboscis (Table S1C). Four NPC2 genes, AAEL012064, AAEL006854, AAEL020314, and AAEL004120 are expressed significantly in the antenna, brain, and proboscis. Although their rank of expression in each tissue varies (Table S2C). While AAEL012064 is expressed the highest in the brain and proboscis, AAEL006854 is the highest expressed gene in the antenna. AAEL001650, the fourth most expressed gene in the brain, shows no expression in the antenna and proboscis (Table S2C). The high expression of these genes in a specific tissue indicates that they may have a significant role in those tissues. In the whole body, AAEL009760 shows a high expression value (Table S2C). Considering that this gene is expressed only the third highest in the midgut and fourth highest in the salivary gland, we speculate that the gene might also be highly expressed in a tissue that has not yet been analysed. Data also indicates that AAEL025019, the potential fatty-acid binding gene, is not expressed in detectable levels in adult females (Table S1A-C). It is unclear whether it is expressed in other tissues, in male mosquitos or in other developmental stages of mosquito life cycle.

### Expression of NPC2 genes post arbovirus infection

Gene expressions in the midgut (Infection initiation site – Midgut infection barrier) and salivary gland (Pathogen transmission site – Salivary gland infection barrier) after an arbovirus infection have been documented [9–12]. We analysed the differential expression of NPC2 genes in uninfected and infected mosquito midgut and salivary gland tissues to understand the potential role of NPC2 proteins during infection. Transcriptome data from the midgut (1-dpi, 4-dpi and 14-dpi for CHIKV, DENV1 or DENV2 with suitable controls, respectively) and salivary gland (7-dpi for CHIKV and 14-dpi for DENV2 or ZIKV) were examined [9–12]. Gene expression data of the brain, antenna and proboscis after an infection, as the roles of these tissues in virus infection have not yet been established.

#### Differential expression in the midgut

Eleven NPC2 genes showed differential expression in the midgut of in uninfected and infected CHIKV mosquitos (Figure 5A). Six genes, AAEL001650, AAEL001654, AAEL006854, AAEL012064, AAEL004120 and AAEL009556 are downregulated (Figure 5A). While AAEL001650, AAEL001654 and AAEL006854 turned down to zero; AAEL012064, AAEL004120 and AAEL009556 are downregulated by 1.9-, 1.7- and 1.4-fold, respectively (Table S3A and S3B). On the other hand, five genes, AAEL009555, AAEL020314, AAEL007591, AAEL015137 and AAEL009760 are upregulated (Figure 5A). AAEL009555 is only expressed post infection. AAEL020314 is upregulated by 2.0-fold, AAEL007591, AAEL009760 and AAEL015137 are each upregulated by about 1.5- fold (Table S3A and S3B).

**Fig 5.**
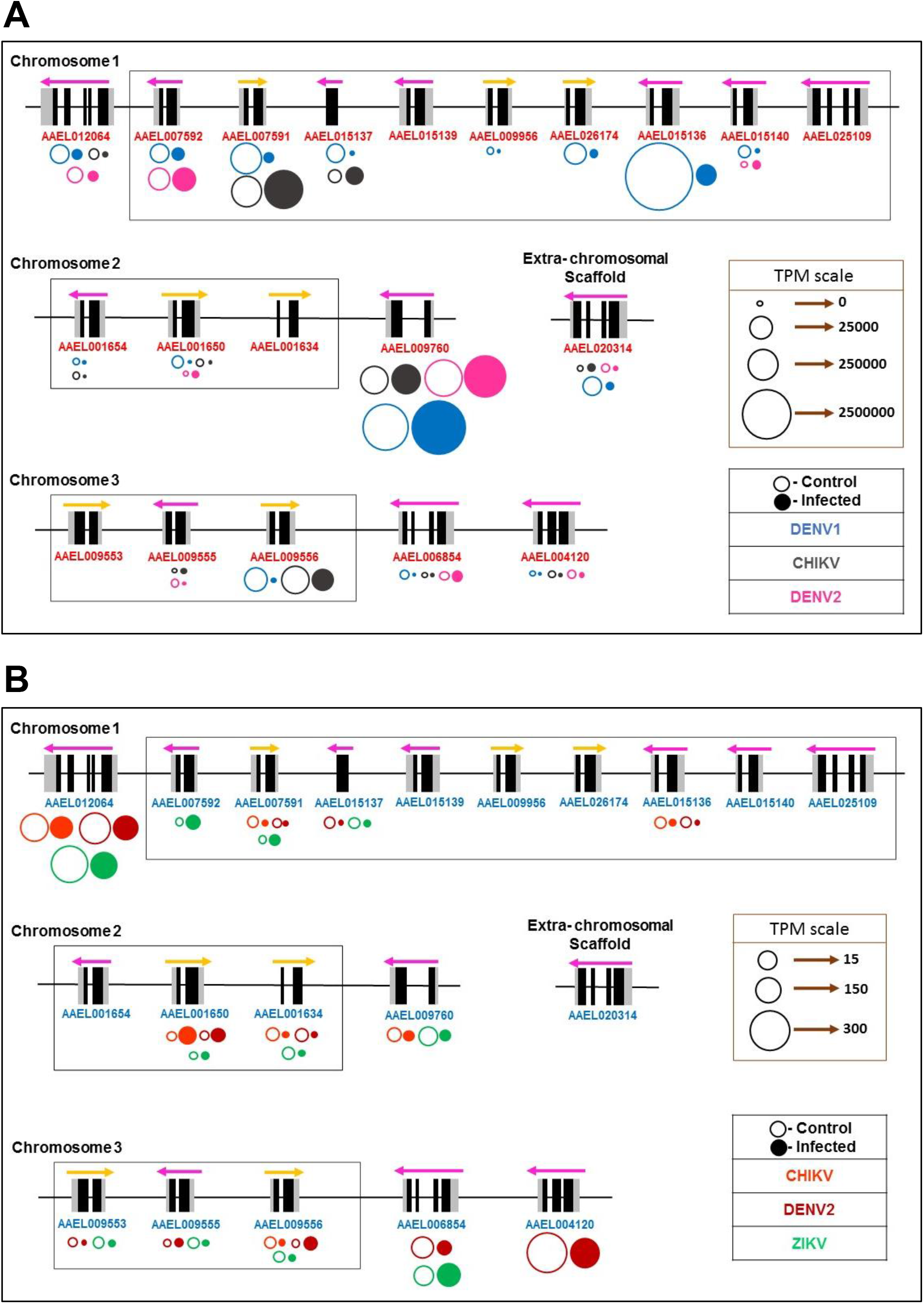
Differential expression of NPC2 genes. **(A)** Differential expression of NPC2 genes in midgut of mosquitos uninfected and infected with CHIKV, DENV1 and DENV2. CHIKV is depicted as grey, DENV1 as blue and DENV2 as pink. **(B)** Differential expression of NPC2 genes in salivary gland of mosquitos uninfected and infected with CHIKV, DENV2 and ZIKV. CHIKV is depicted as orange, DENV2 as maroon and ZIKV as green. The area of circles represents the expression level. The expression in uninfected (control) is depicted with open circles and infected with closed circles.

We analysed the differential expression of NPC2 genes in uninfected and infected mosquitos, infected with two different serotypes of DENV – DENV1 and DENV2 [9], [10]. In the case of DENV1 at 4-dpi in the Thailand mosquito strain, 14 genes are downregulated (Figure 5A). The expression of AAEL001650, AAEL001654 and AAEL009956 is fully turned off post DENV1 infection (Table S3C and S3D). The order of differential expression for eleven genes along with their fold change is AAEL007591 (76.3x) > AAEL015137 (75.7x) >> AAEL015136 (54.7x) >> AAEL015140 (24.5x) > AAEL009556 (22.4x) > AAEL006854 (14.3x) > AAEL004120 (10.7x) > AAEL026174 (9x) > AAEL020314 (4.6x) > AAEL012064 (4x) >> AAEL007592 (1.5x) (Table S3C and S3D). On the other hand, AAEL009760 (14.6x) is the only gene that is upregulated post-DENV1 (Table S3D).

In the case of DENV2 at 14-dpi in the Chetumal mosquito strain, five genes are downregulated (Figure 5A). The expression of AAEL009555 is turned off completely post-DENV2 infection (Table S3E). The order of differential expression for four genes along with their fold change is AAEL020314 (7x) >> AAEL012064 (2.4x) > AAEL004120 (1.9x) > AAEL009556 (1.2x) (Table S3E and S3F). On the other hand, four genes are upregulated in the order AAEL015140 (2.8x) > AAEL001650 (1.8x) > AAEL006854 (1.6x) > AAEL009760 (1.2x) (Table S3E and S3F). Independent studies also show the upregulation (2.2x) of AAEL001650 and AAEL006854 was observed 7-dpi in DENV2 infected midgut of Rockefeller (laboratory-maintained) and Caribbean (field-derived) strains of mosquitos [19]. Although both genes were silenced in these strains of mosquitos, only AAEL001650 showed a significant decrease of median DENV2 titre in the midgut, while silencing of AAEL006854 had no significant effect on infection [19]. In a recent study, knockdown of gene AAEL015136 lead to a decrease in DENV2 infection in the Cali-S susceptible strains, whereas it increased the infection rate in the Cali-MIB refractory strains [36]. The authors concluded that such contradictory observation is probably due to distinct roles played by AAEL015136 in different mosquito strains [36]. This gene did not show any significant differential expression in Chetumal strain of mosquitos (Figure 5A and Table S3E).

NPC2 genes AAEL004120, AAEL012064 and AAEL009556 are downregulated post all three infections (Figure 5A). Whereas, AAEL009760 is upregulated post all three infections. Four genes, AAEL004120, AAEL020314, AAEL012064 and AAEL009556 are downregulated in both DENV1 and DENV2. Whereas, AAEL001650 and AAEL006854 are both downregulated in DENV1 and CHIKV (Figure 5A). Interestingly, AAEL001650, AAEL015140 and AAEL006854 showed contrasting results in midgut infected with DENV1 and DENV2. They are downregulated in DENV1 and upregulated in DENV2 (Figure 5A). These observed differential expression profiles of NPC2 genes may be due to either viral serotypes (DENV1 vs. DENV2), distinct mosquito strains (Thailand vs. Chetumal) or even different time points post-infection (4-dpi vs. 14-dpi). Overall, the above analysis on the differential expression in the midgut showed that NPC2 genes are more downregulated than upregulated in midgut infected with CHIKV, DENV1 and DENV2 (Figure 5A).

#### Differential expression in the salivary gland

In CHIKV infected salivary gland at 7-dpi, six genes, AAEL012064, AAEL001634, AAEL009760, AAEL006854, AAEL004120 and AAEL007591 are downregulated (Figure 5B). The expression of AAEL001634 and AAEL007591 are turned off post-CHIKV (Table S4A and S4B). The order of gene expression of rest of the genes along with the fold change is AAEL009760 (1.4x) > AAEL012064 (1.3x) > AAEL006854 (1.1x) > AAEL004120 (1.0x). AAEL001650 (12.3x) is the only gene upregulated (Table S4A and S4B).

In the case of DENV2 at 14-dpi, eight genes are downregulated (Figure 5B). AAEL007591, AAEL009553, AAEL015136 and AAEL015137 are turned off in DENV2 infected salivary gland (Table S4C and S4D). The order of gene expression of other genes along with the fold change is AAEL001634 (3.5x) > AAEL006854 (2.8x) > AAEL004120 (1.7x) > AAEL012064 (1.5x). On the other hand, three genes are upregulated AAEL009556 (6.4x) > AAEL001650 (5.1x) >> AAEL009555 (1.3x) (Table S4C and S4D).

In ZIKV-infected salivary gland at 14-dpi, seven genes are downregulated (Figure 5B). AAEL009553, AAEL009555, AAEL009556 and AAEL015137 are turned off completely post-ZIKV infection (Table S4E and S4F). AAEL009760 (2.6x) > AAEL012064 (1.6x) > AAEL001634 (1.4x) is the order of gene expression for the other downregulated genes. On the other hand, three genes are upregulated in the order AAEL007591 (2.6x) > AAEL006854 (1.5x) > AAEL001650 (1.3x). AAEL007592 is expressed only post-ZIKV infection (Table S4E and S4F).

Seven genes (AEL001654, AAEL009956, AAEL015139, AAEL015140, AAEL020314, AAEL026174 and AAEL025109) are not expressed in the uninfected or infected (CHIKV, DENV2 or ZIKV) salivary gland. Interestingly, in the salivary gland infected with CHIKV, DENV2 or ZIKV, AAEL012064 and AAEL001634 are downregulated and AAEL01650 is upregulated (Figure 5B). As these differential expression profiles are derived from the same strain of mosquitos (Singapore), we speculate that these three genes are important in replication of arboviruses in the salivary gland. On the other hand, AAEL006854 and AAEL007591 are downregulated in salivary gland infected with CHIKV and DENV2, but are upregulated when infected with ZIKV (Figure 5B). The contrasting expression profiles may be due to different roles played by AAEL006854 and AAEL007591 in the infection process of CHIKV, DENV2 or ZIKV. Similar to midgut, most genes were downregulated rather than upregulated in the infected salivary gland (Figure 5B). AAEL012064 is the only gene downregulated in both midgut and salivary gland tissues in all virus-infected mosquitos. It will be interesting to understand the functional role of this protein in arbovirus infections.

### Transcription factors (TFs) regulating NPC2 gene expression

The binding of specific TFs at the PPR sites (*cis* elements) may regulate the expression of NPC2 genes. The variation of expression of closely related genes are regulated by variation in *cis* elements and/or expression of TFs themselves. We focused on two groups of *A. aegypti* NPC2 genes whose members share high identity and encode potential cholesterol-binding proteins (Figure 1B, i and ii). Group A consists of two genes AAEL006854 (chromosome 3) and AAEL020314 (extra-chromosomal scaffold) that share the highest identity (protein 99.24%) (Figure 1B(i)). Group B consists of three genes AAEL009553, AAEL009555, AAEL009556 that form a cluster in chromosome 3 and show high identity (protein 73.46-91.36%) to each other (Figure 1B(ii)). Only AAEL009555, due to I126T substitution may not bind cholesterol (Figure 3C). We analysed the expression patterns of these two groups and attempted to identify the TFs that bind to the PPR and may play a role in their differential expression.

#### Differential expression of group A genes and TFs regulating their expression

In uninfected mosquitos, AAEL006854 and AAEL020314 are expressed in the antenna, brain, proboscis, and midgut (Table S1C). While AAEL006854 is expressed in the midgut and salivary gland of uninfected mosquitos, AAEL020314 is expressed only in the midgut.

The binding sites for the TFs in the PPR of both these genes were identified (Figure 6A). As expected, the *cis* elements are similar to each other, and they shared many common and overlapping sites. The TFs expressed exclusively in the midgut and salivary gland were plotted separately (Figure 6, B and C). We identified two and one unique TF-binding sites in AAEL006854 and AAEL020314, respectively. Two TFs – NF-kB and TBP are unique to AAEL006854 (Figure 6A) and are expressed in the midgut and salivary gland (Table S5A and S6A). A TF-Hunchback (Hb; AAEL000894), unique to AAEL020314, is found -400 bp from the transcription initiation site (Figure 6A). Interestingly, Hb is not expressed in either midgut or salivary gland (Table S5A and S6A).

**Fig 6.**
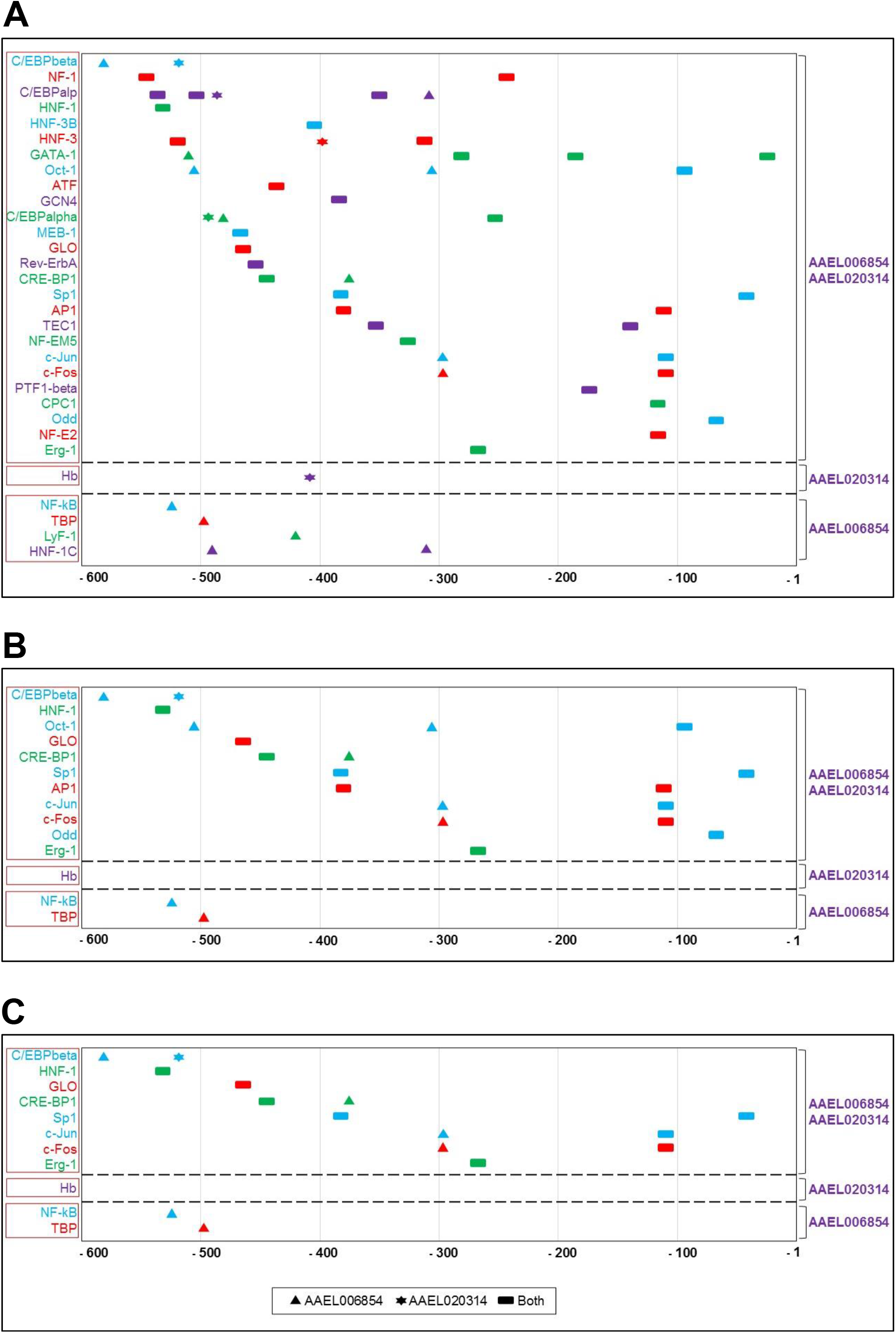
TF binding sites (*cis* elements) and TF in the expression of group A genes. **(A)** As AAEL006854 and AAEL020314 genes share high identity, they share many overlapping sites. The top segment depicts the TFs that are common to both the genes, the second and third segments are TFs unique to AAEL020314 and AAEL006854, respectively. **(B)** The TFs of group A genes that are expressed in the midgut. **(C)** The TFs of group A genes that are expressed in the salivary glands. The TF binding sites of PPR of group A genes were predicted by TRANSFAC 4.0 and AliBaba 2. TF sites are depicted for AAEL006854, ▲; AAEL020314, ✶; and overlapping sites, ▄. The colour of the symbol matches that of the TF name.

In the midgut of infected mosquitos, AAEL006854 and AAEL020314 are downregulated post-DENV1 infection (Figures 7A and 8A). The two genes show an opposite expression pattern post-DENV2 infection; AAEL006854 is upregulated and AAEL020314 is downregulated (Figures 7A and 8A). Interestingly, six TFs (CRE-BP1, AP1, c-Jun, c-Fos, Odd and NF-kB) are significantly upregulated post-DENV1 infection but downregulated post-DENV2 infection (Figure 8B and Table S5B). When these six TFs are downregulated post-DENV1 infection, AAEL006854 is upregulated. Whereas when the six TFs are upregulated post DENV-2 infection, AAEL006854 is downregulated (Table S5B). Interestingly, the same pattern was not observed in AAEL020314. Hence, we speculate that these TFs may play an opposing role in the differential expression of AAEL006854 in midgut infected with DENV1 or DENV2.

**Fig 7.**
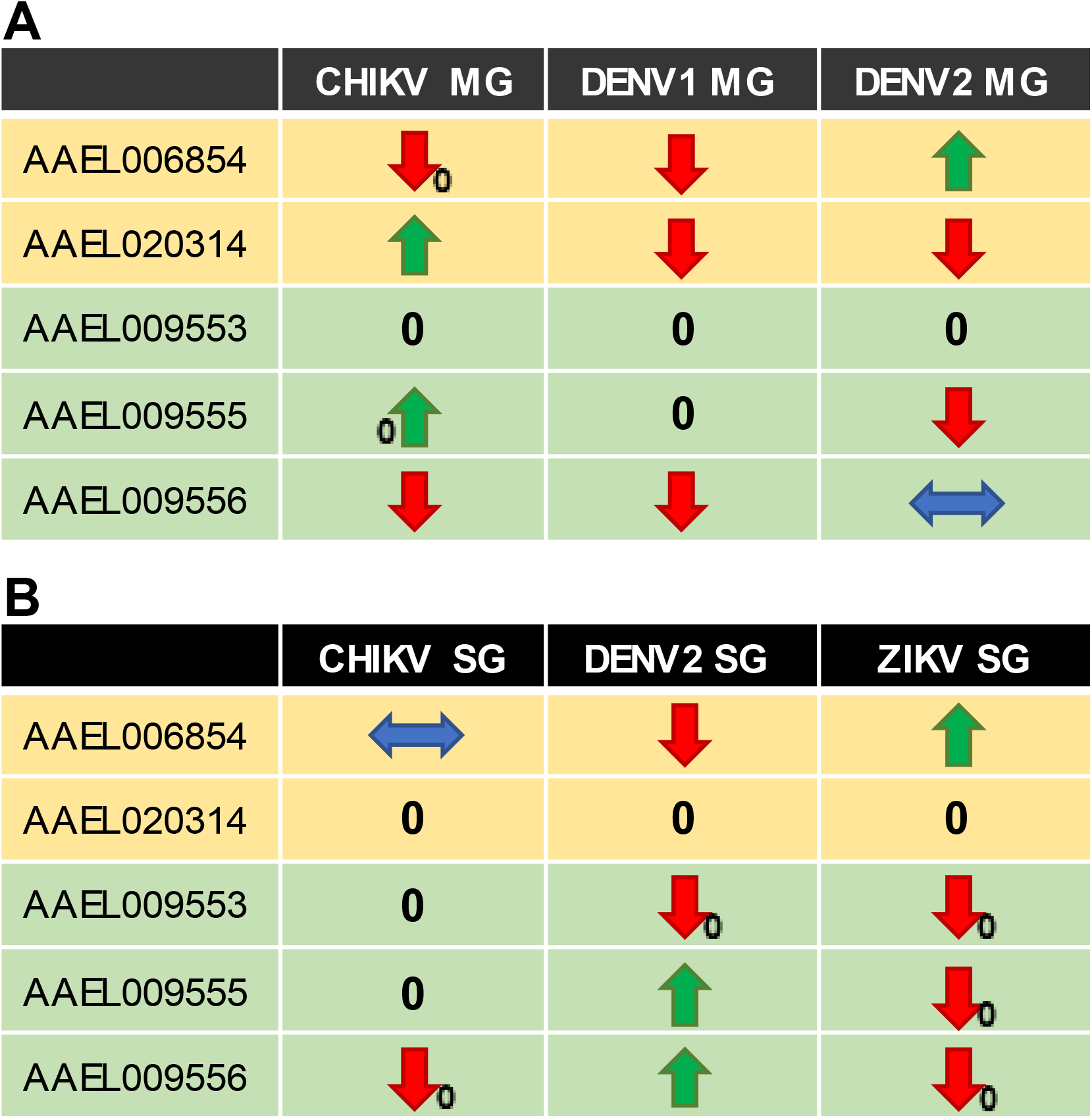
Differential expression of groups A and B NPC2 genes in virus-infected mosquitos. **(A)** Differential expression in the midgut of mosquitos infected with CHIKV, DENV1 or DENV2. **(B)** Differential expression in the salivary gland of mosquitos infected with CHIKV, DENV2 or ZIKV. The differential expression of group A (orange), and group B (green). 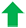 , upregulation; 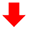 , downregulation; 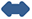, no differential expression post-infection. A ‘0’ in the subscript next to an arrow means that the gene is turned off post infection or expressed only post infection. ‘0’, indicates no detectable expression.

**Fig 8.**
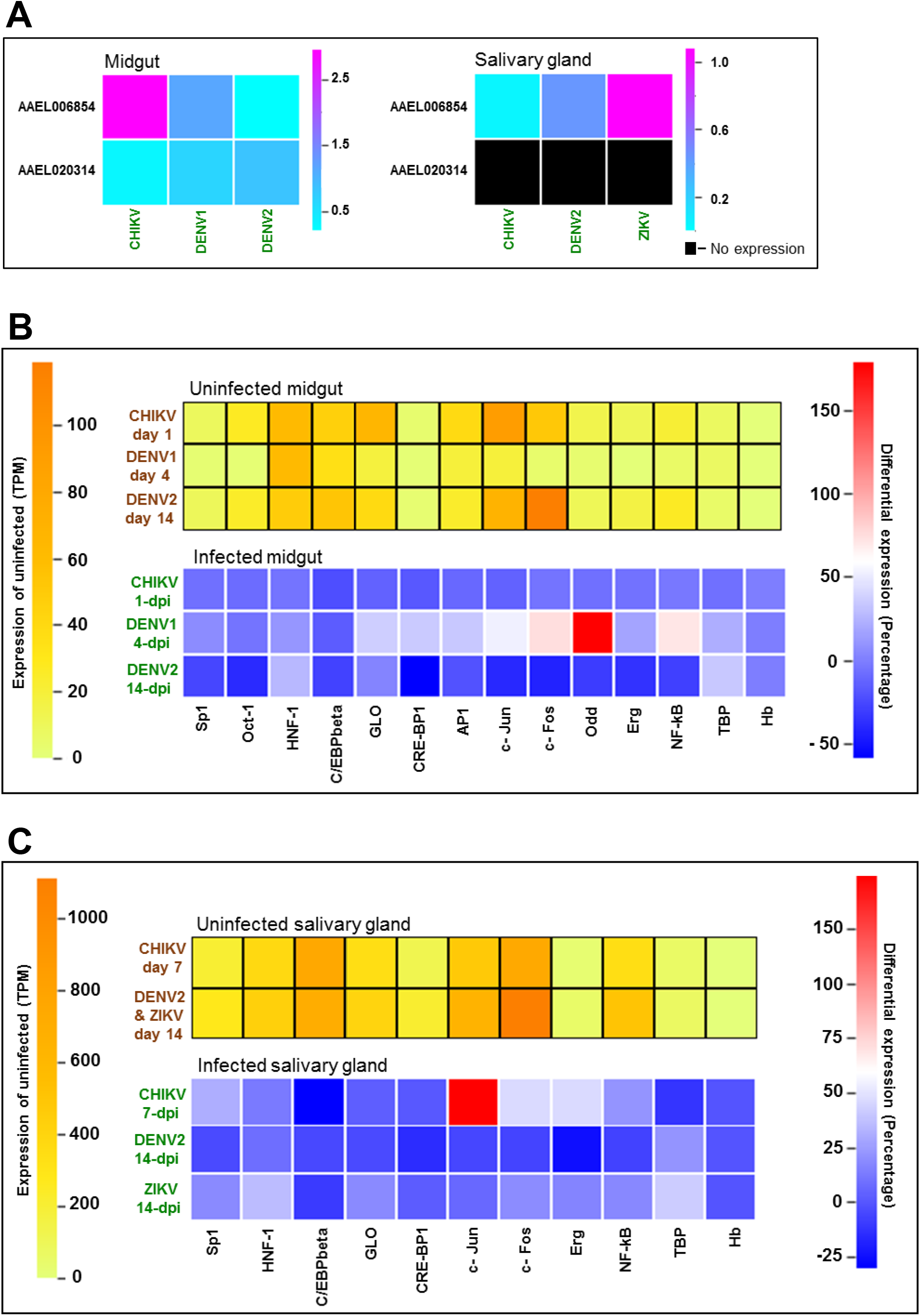
Differential expression of group A NPC2 and relevant TFs in virus-infected mosquitos. **(A)** The differential expression of AAEL006854 and AAEL020314 in (a) midgut infected with CHIKV, DENV1 or DENV2, and (b) salivary gland infected with CHIKV, DENV2 or ZIKV. The change in expression is represented in log10 of the fold. The black boxes represent no detectable gene expression in the tissue. **(B)** The heatmaps represent TF expression in the uninfected midgut (top, represented in TPM) and differential expression in midgut infected with CHIKV, DENV1 or DENV2 (bottom, represented in percentage). **(C)** The heatmaps represent TF expression in the uninfected salivary gland (top, represented in TPM) and differential expression in the salivary gland infected with CHIKV, DENV2 and ZIKV (bottom, represented in percentage).

In the salivary gland, AAEL006854 is downregulated post-DENV2 infection (Figure 7B), while it is upregulated in ZIKV infection and shows no significant change post-CHIKV infection (Figure 7B). TBP is upregulated post-DENV2 and -ZIKV infection (Figure 8C and Table S5C), while its expression remains unchanged post-CHIKV infection (Figure 8C and Table S5C). On the other hand, NF-kB is upregulated post-CHIKV infection, but no significant change is seen post-DENV2 and -ZIKV infections (Figure 8C and Table S5C). Interestingly, three TFs (c-Jun, c-Fos and Erg) are significantly upregulated in CHIKV infected salivary gland.

#### Differential expression of group B genes and TFs regulating their expression

AAEL009553, AAEL009555, and AAEL009556 displayed different expression patterns in uninfected mosquitos (Table S1-S4). In the uninfected mosquitos, AAEL009553 is expressed in small amounts only in the antenna and salivary gland (Table S1). AAEL009555 is expressed in small amounts only in the salivary gland and midgut (Table S1). Whereas, AAEL009556 is expressed significantly in the midgut, and in small amounts in the brain and salivary gland (Table S1).

The *cis* elements in the PPR of group B genes were identified (Figure 9A). The TFs expressed exclusively in the midgut or salivary gland were plotted separately (Figure 9B and 9C). AAEL009553 shows no expression in the midgut of uninfected and infected (CHIKV, DENV1 and DENV2) mosquitos (Figures 7A and 10A). AAEL009555 shows no expression in the midgut of uninfected and infected (DENV1) mosquitos (Figures 7A and 10A). Five TFs common to these two genes post-DENV1 infection are RAP1, NF-kB, YY1, MCM1 and TBP (Figure 9B). Additionally, AAEL009553 has two unique *cis* elements for Pit-1a and GLO and AAEL009555 has three unique *cis* elements for CRE-BP1, SGF, and GABP (Figure 9B). All these TFs are upregulated post-DENV1 infection (Figure 10B and Table S7B) and might play a role in the expression of AAEL009553 and AAEL009555.

**Fig 9.**
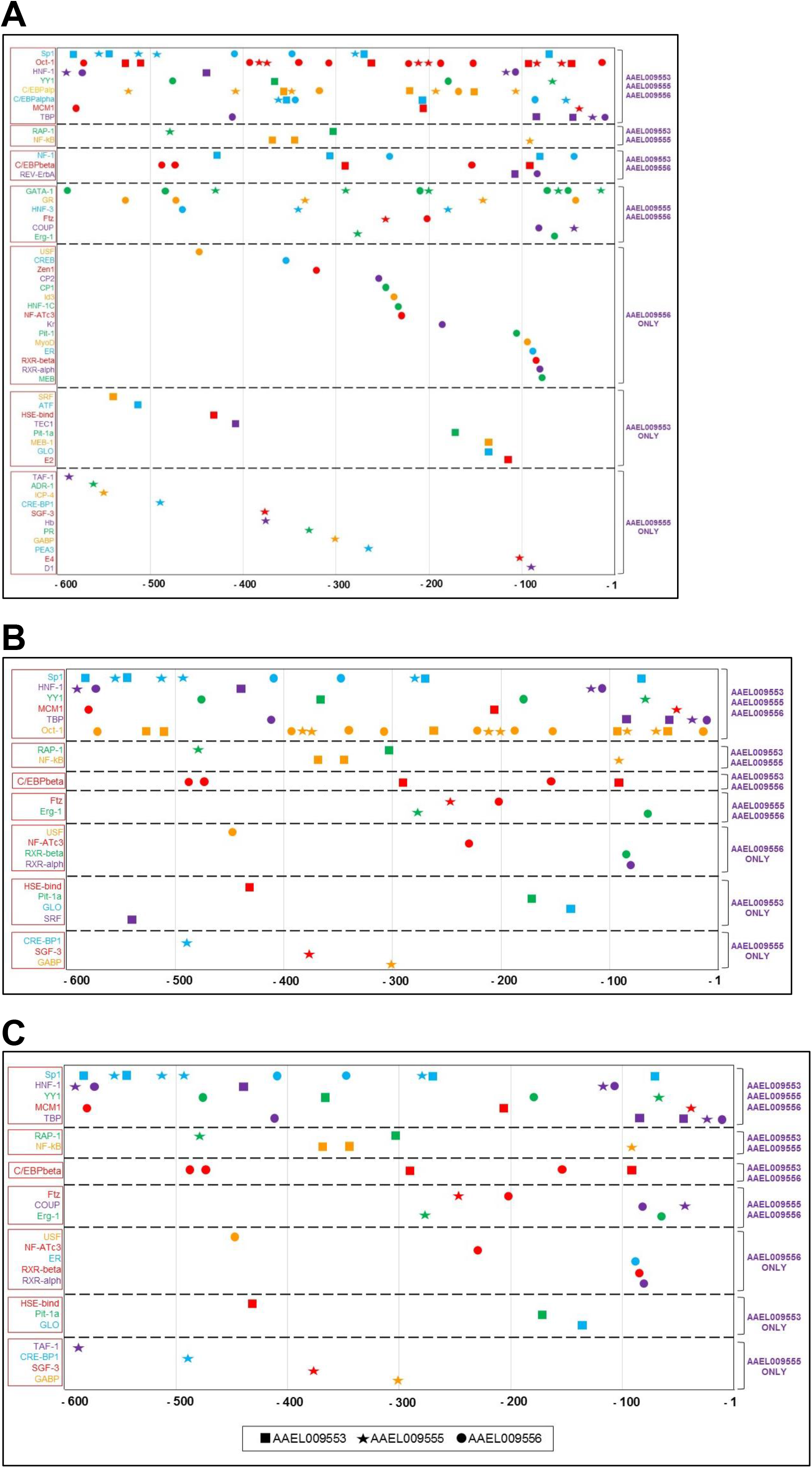
TF binding sites (*cis* elements) and TF in the expression of group B genes. **(A)** The top segment depicts the TFs that are common to all the genes, the second, third and fourth segments are TFs common to any two genes, and the fifth, sixth and seventh segments are TFs unique to each gene, respectively. **(B)** The TFs of group B genes that are expressed in the midgut. **(C)** The TFs of group B genes that are expressed in the salivary glands. The TF binding sites of PPR of group B genes were predicted by TRANSFAC 4.0 and AliBaba 2. TF sites are depicted for AAEL009553, ■ ; AAEL009555, ★; and AAEL009556, ● . The colour of the symbol matches that of the TF name.

**Fig 10.**
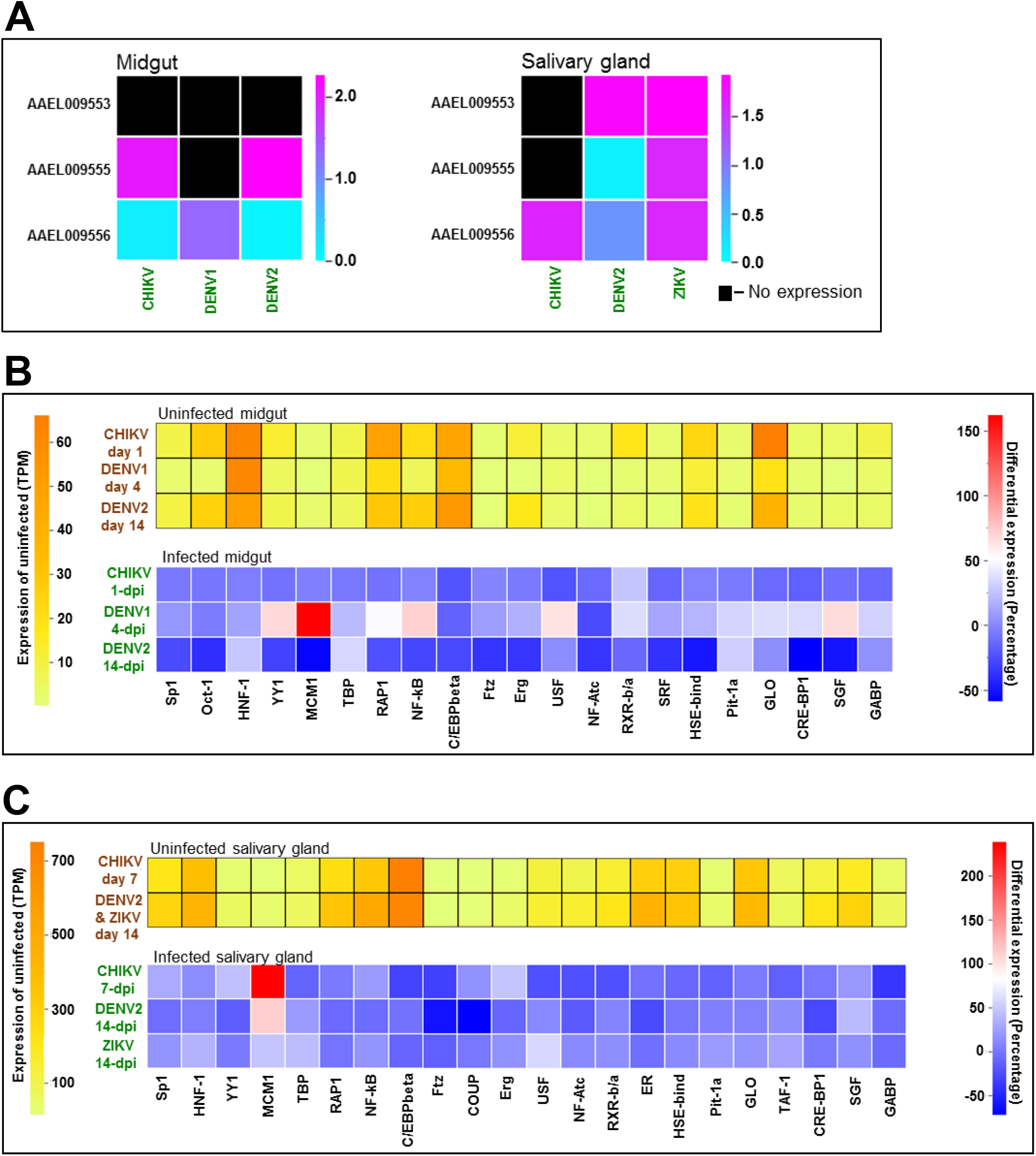
Differential expression of group B NPC2 and relevant TFs in virus-infected mosquitos. **(A)** The differential expression of AAEL009553, AAEL009555 and AAEL009556 in (a) midgut infected with CHIKV, DENV1 or DENV2, and (b) salivary gland infected with CHIKV, DENV2 or ZIKV. The change in expression is represented in log10 of the fold. The black boxes represent no detectable gene expression in the tissue. **(B)** The heatmaps represent TF expression in the uninfected midgut (top, represented in TPM) and differential expression in midgut infected with CHIKV, DENV1 or DENV2 (bottom, represented in percentage). **(C)** The heatmaps represent TF expression in the uninfected salivary gland (top, represented in TPM) and differential expression in the salivary gland infected with CHIKV, DENV2 and ZIKV (bottom, represented in percentage).

Further, AAEL009556 is downregulated in the midgut of mosquitos infected with CHIKV and DENV1 (Figures 7A and 10A). TF RXR-beta/alpha is commonly upregulated post CHIKV and DENV1 infection (Figure 10B and Table S7B). USP, the drosophila homolog of RXR, acts as a repressor in the drosophila eye development [37]. Additionally, TFs (USF and NF-Atc) unique to AAEL009556 are downregulated in CHIKV and DENV1 infections, respectively (Figure 10B and Table S7B). Interestingly, USF is known to activate the hRFC promoter, while NF-Atc is known to activate the HIV replication in T cells [38], [39]. These unique TFs along with RXR-beta/alpha could play a role in the downregulation of AAEL009556 in midgut infected with CHIKV and DENV1.

In the salivary gland, group B genes are downregulated post-ZIKV infection (Figure 7B). The TFs common to the three PPRs, HNF-1, MCM1 and TBP, are upregulated significantly post-ZIKV infection (Figure 10C and Table S7B). TFs namely, Pit1a, TAF1 and USF, are unique to AAEL009553, AAEL009555 and AAEL009556, respectively (Figure 9C). All these TFs are upregulated significantly in ZIKV infected salivary gland (Figure 10C and Table S7B). All the above TFs might collectively play a role in the downregulation of group B genes in ZIKV infected salivary gland.

AAEL009555 and AAEL009556 are upregulated in the salivary gland infected with DENV2 (Figure 7B). The TFs common for the two genes are Ftz and COUP. Ftz and COUP are downregulated post-DENV2 infection (Figure 10C and Table S8B). The overexpression of Ftz is reported to activate the E75A gene [40], while COUP plays dual roles of an enhancer and a suppressor [41–43]. We speculate that the differential expression of these two TFs mainly play a role in the upregulation of AAEL009555 and AAEL009556 post-DENV2.

AAEL009556 is downregulated post-CHIKV infection in the salivary gland (Figure 8B). This gene has two unique TF binding sites – USF and RXR beta/alpha which are downregulated in CHIKV infected salivary gland (Figure 10C and Table S8B). We speculate that these TFs potentially play a role in the downregulation of AAEL009556 in CHIKV infected salivary gland, similar to CHIKV infected midgut.

## Discussion

In mammals, only one highly conserved, ubiquitously expressed gene encodes for NPC2 protein, which participates in cholesterol binding and transport in all cells [28]. By contrast, arthropods express several NPC2 proteins, whose functions are unclear. In *A. aegypti*, the NPC2 superfamily contains 20 proteins that are diverse and expressed in the midgut, salivary gland, brain, antenna, and proboscis. However, none of them appears to be expressed ubiquitously. These NPC2 proteins are poorly characterized, and no information on the function and structure-function relationships are available. A comparison of *A. aegypti* NPC2 proteins with bovine NPC2 protein and its residues involved in binding cholesterol helped us understand the potential interaction of mosquito NPC2 proteins with cholesterol or related sterols (Figure 3C). Several *A. aegypti* NPC2 proteins have unfavourable mutations that would disrupt the ligand binding or potentially modify the conformation of the protein, resulting in the loss of ligand binding (Figure 3C). Based on our analysis, we identified four NPC2 proteins (AAEL006854, AAEL020314, AAEL009553 and AAEL009556) as potential cholesterol-binding proteins (Figure 3C). Most residues of NPC2 interacting with human NPC1 for cholesterol transfer are altered in all four potential cholesterol-binding proteins (Figure 3D). Similarly, *A. aegypti* NPC1 also have non-conserved mutations replacing the residues of the human NPC1 functional site (Figure S4). Hence, it is unclear whether NPC1 and NPC2 proteins in the mosquito are involved in cholesterol transfer or not. However, their roles could be different from that of vertebrate NPC2 protein.

The potential cholesterol-binding NPC2 proteins have a varied expression in the uninfected tissues of *A. aegypti*. While AAEL006854 and AAEL020314 are expressed highly in the antenna, brain, and proboscis, AAEL009553 and AAEL009556 are expressed in small amounts only in the antenna and brain, respectively (Table S1C). AAEL006854 and AAEL009556 are expressed in the midgut and salivary gland, whereas AAEL020314 and AAEL009553 are expressed only in the midgut and salivary gland, respectively (Figure 7, A and B). The varied expression of the potential cholesterol-binding proteins in different mosquito tissues may indicate different roles played by them in these tissues. Interaction of these NPC2 proteins with various sterols as well as with *A. aegypti* NPC1 proteins may help delineate their physiological functions. Studies by Jupatanakul et al. showed that AAEL006854 (a NPC2) and AAEL009531 (a NPC1) act as agonist of DENV1 infection in the midgut [19]. Since none of the *A. aegypti* NPC2 genes are ubiquitously expressed, it would be interesting to understand the players and the mechanism of intracellular cholesterol transport in invertebrates.

Studies suggest that the NPC2 protein family in arthropods are involved in chemical communication by binding to certain semiochemicals [29], [35]. NPC2 of the worker ant *Camponotus japonicus* binds to oleic acid, a long-chain fatty acid proposed to aid in chemical communication [29]. We compared mosquito NPC2 proteins to the residues of the binding pocket of ant NPC2 (Figure 4C). Most of the mosquito proteins presented an unfavourable environment for fatty acid binding. Only AAEL025109 comprised conserved residues for fatty acid binding (Figure 4C). As a result, we predicted that this protein may bind to long-chain fatty acid or alcohol. Unlike ant NPC2, AAEL025109 is not expressed in the antenna and thus, may have a distinct physiological role. Whereas the highly expressed genes in the antenna (AAEL006854, AAEL020314, AAEL004120 and AAEL012064) cannot bind to a long-chain fatty acid due to various mutations in the binding residues, especially the mutation of the double lysine site that form the hydrogen bonds with the fatty acid. Further, it is not clear whether AAEL025109 is expressed in the antenna or other tissues of male mosquitos or in the egg, larval or pupal stages. Tissue-specific expression and interaction with ligands may help deciphering AAEL025109’s physiological role.

*A. aegypti* is the primary or secondary vector for arbovirus infections like DENV, CHIKV, and ZIKV. The transmission takes place during the blood meals when a mosquito injects its saliva into the host to combat host responses and maintain continuous blood flow [8]. Several mosquito salivary proteins aid in this process and actively contribute to enhanced transmission of viruses [8]. Several studies evaluated transcriptomics of uninfected and virus-infected mosquitos [9–14]. To understand the role of NPC2 genes in infected mosquitos, we focussed only on their differential expression in the midgut and salivary gland, two key tissues that play critical role in pathogen transmission. We also analysed their expression in other tissues including antenna, brain and proboscis isolated from uninfected mosquitos. Four genes (AAEL012064, AAEL004120, AAEL006854 and AAEL020314) are highly expressed in the antenna, brain, and proboscis (Table S2C). Whereas ten genes (AAEL001634, AAEL007591, AAEL009555, AAEL009956, AAEL015136, AAEL015137, AAEL015139, AAEL015140, AAEL026174 and AAEL025109) show no expression in these tissues (Table S1C).

In the midgut infected with CHIKV, DENV1 or DENV2, while AAEL004120, AAEL012064 and AAEL009556 are downregulated, AAEL009760 is upregulated (Figure 5A). In a previous study, AAEL004120 exhibited no differential expression post-DENV2 infection in any tissue compartments, suggesting that it may not have a role in DENV infection [19]. However, as shown here, the transcriptomic data from other studies show that this gene is downregulated in midgut infected with CHIKV (1.7x), DENV1 (10.7x) or DENV2 (1.9x). The prevalent differential expression of AAEL004120 in distinct mosquito strains (Puerto Rico vs. Thailand vs. Chetumal) and arboviruses (CHIKV, DENV1 and DENV2) indicates a potential role of this gene in these infections. Further, three genes (AAEL001650, AAEL015140 and AAEL006854) are downregulated post-DENV1 and upregulated post-DENV2 infections (Figure 5A). These contrasting differential expression profiles of NPC2 genes may be due to (a) different virus serotypes (DENV1 vs. DENV2); (b) distinct mosquito strains (Thailand vs. Chetumal); and/or (c) different dissection time points post-infection (4-dpi vs. 14-dpi). In the salivary gland infected with CHIKV, DENV2 or ZIKV, while AAEL001634 and AAEL012064 are downregulated, AAEL001650 is upregulated (Figure 5B). As these differential profiles were observed in the same strain of mosquito (Singapore), these genes could potentially have an important role in arbovirus infections in the salivary gland. Overall, we observed that most genes were downregulated rather than upregulated post an infection in both the midgut and salivary gland.

As *cis* elements and TFs are responsible for expression regulation, we evaluated the PPR of two groups of closely related genes encoding potential cholesterol-binding proteins. We also analysed the differential expression of relevant TFs in the midgut and salivary gland. Six TFs that are involved in the expression regulation of AAEL006854, CRE-BP1, AP1, c-Jun, c-Fos, Odd and NF-kB are significantly upregulated or downregulated, respectively, post-DENV1 or -DENV2 infection in the midgut. This disparity in expression, at the first glance, appears to be due to different serotypes of DENV. In fact, such differences could be due to various other factors (discussed below). Despite the limitations, it is clear that specific TFs indeed would play key roles in the differential expression of AAEL006854 within each set of experiments. Interestingly, no such pattern was found in AAEL020314. A TF unique to AAEL020314, Hb (AAEL000894), is not expressed in either uninfected or infected midgut and salivary gland. Since AAEL020314 is expressed in the midgut, it is hard to draw a conclusion on the role of Hb in the expression of AAEL020314. Although the potential role of Hb may not be ignored. Oct-1, a TF binding to *cis* elements of groups A and B genes, is exclusively expressed in the midgut (Table S5A and S7A). Similarly, in group B genes, TFs COUP (common to AAEL009555 and AAEL009556) and TAF1 (unique to AAEL009555) are expressed only in the salivary gland (Table S8A). Knockdown experiments of these TFs in the specific tissue of group A and group B genes would help in understanding their tissue- and gene-specific roles.

The above conclusions should be considered with the following considerations. The transcriptomic data was collected from different strains of mosquitos, timepoints (dpi) and sequencing methods. Further, the PPR data of *cis* elements was analysed from the only genome of Liverpool strain that is publicly accessible. These limitations could be overcome by performing the genomic and transcriptomic studies in the same strain of mosquito with appropriate controls for all the arbovirus infections. Knockdown experiments of specific genes may further help in understanding the role of NPC2 genes in virus transmission.

