## Supplementary Information for "Nieman-Pick Type C2 proteins in *Aedes aegypti*: Their Structure-Function relationships and Expression in Uninfected *versus* Virus-infected Mosquitos"

#### Supplementary Figures

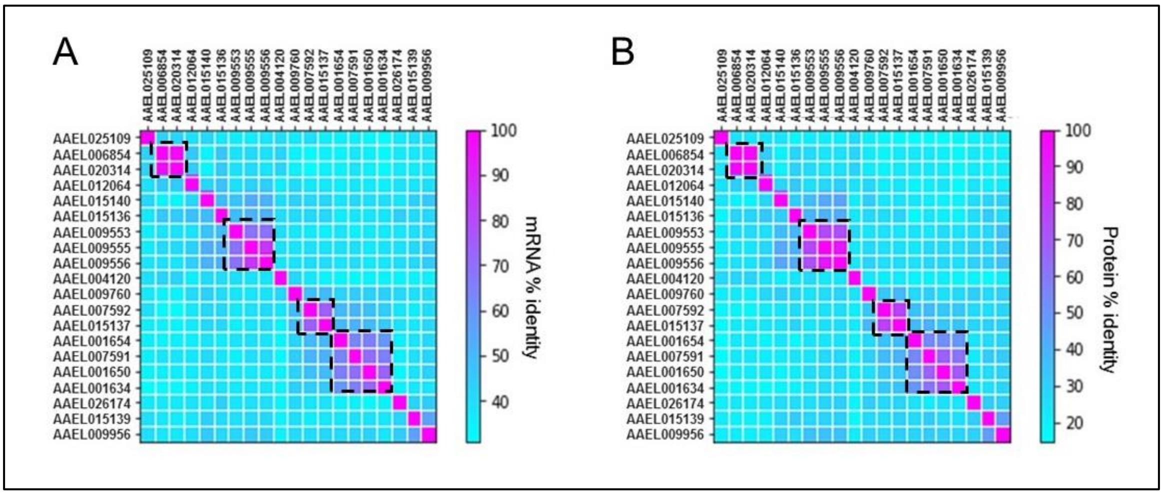

**Fig S1. Sequence identity of *A. aegypti* NPC2.** The heatmaps depict the mRNA (A) and protein (B) identity (%) of 20 *A. aegypti* NPC2. Groups A (AAEL006854 and AAEL020314), B (AAEL009553, AAEL009555 and AAEL009556), C (AAEL001654, AAEL007591, AAEL001650, and AAEL001634) and D (AAEL007592 and AAEL015137) form clusters of high identity on the heatmap (marked with black dotted boxes).

| | AAEL007592 | AAEL007591 | AAEL015137 | AAEL015139 | AAEL009956 | AAEL026174 | AAEL015136 | AAEL015140 | AAEL025109 |
|---|---|---|---|---|---|---|---|---|---|
| AAEL007592 | | 37.33 | 77.63 | 25.5 | 29.05 | 28.08 | 26.53 | 21.48 | 15.15 |
| AAEL007591 | 51.43 | | 41.06 | 30.41 | 32.65 | 21.92 | 24.49 | 22.97 | 13.64 |
| AAEL015137 | 73.64 | 52.85 | | 29.14 | 30.67 | 29.25 | 29.05 | 23.84 | 15.04 |
| AAEL015139 | 39.96 | 42.67 | 40.96 | | 47.33 | 30.61 | 30.61 | 28.00 | 21.97 |
| AAEL009956 | 41.03 | 44.7 | 44.25 | 56.07 | | 30.14 | 25.85 | 30.00 | 18.18 |
| AAEL026174 | 36.05 | 34.85 | 36.91 | 38.02 | 41.69 | | 26.21 | 26.71 | 18.37 |
| AAEL015136 | 40.33 | 38.21 | 41.22 | 41.78 | 45.11 | 45.66 | | 31.76 | 17.56 |
| AAEL015140 | 40.32 | 39.22 | 40.67 | 38.48 | 41.47 | 39.69 | 49.66 | | 23.13 |
| AAEL025109 | 39.02 | 35.51 | 38.05 | 40.23 | 39.76 | 37.03 | 35.32 | 40.55 | |

**Fig S2. Sequence identity among nine NPC2 genes forming a large cluster in chromosome 1.** The sequence identity of nine genes that are located in chromosome 1 and form a cluster**.** The lowest percentage identity in mRNA is about 35%, whereas it is about 14% in proteins. NPC2 expressing low identity are marked in green, whereas the high percentages are marked in purple.

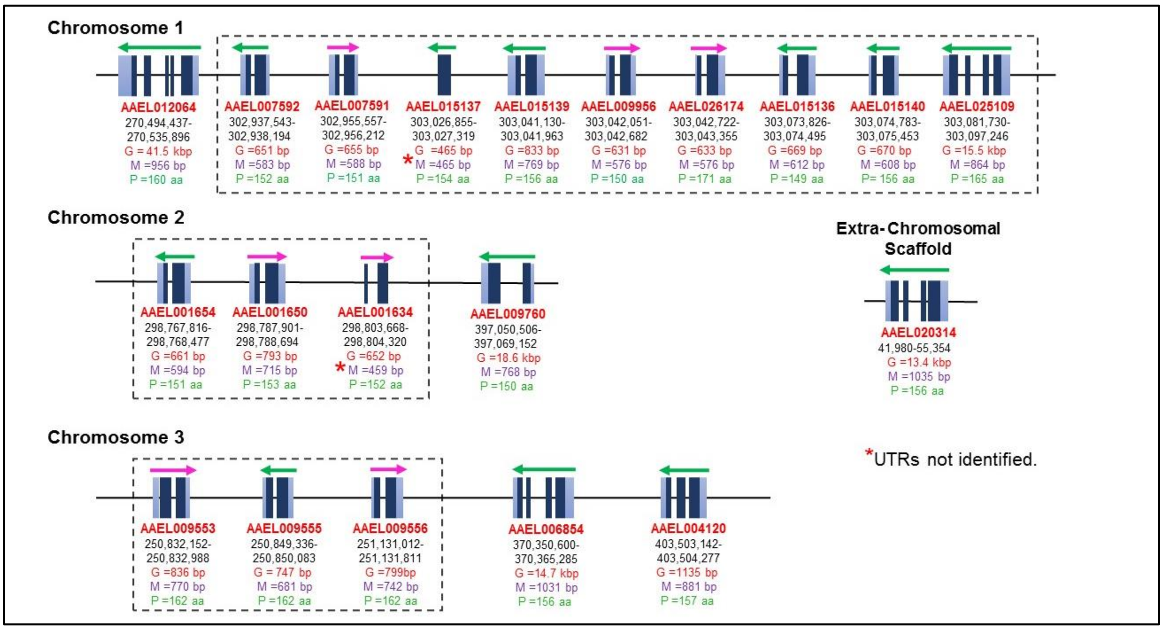

**Fig S3. Intron-exon profile, orientation, location and sizes of gene, mRNA and full-length protein of *A. aegypti* NPC2 genes.** Intron-exon profiles for each gene is indicated with blue boxes, exons (dark blue boxes), introns (gaps between boxes), and 5’ and 3’ UTRs (light blue boxes). The UTRs of some genes (*) have not been identified. The chromosome location, and sizes of gene (G), mRNA (M) and protein (P) are indicated in appropriate units of base pairs and amino acids. The direction of transcription is shown above each gene.

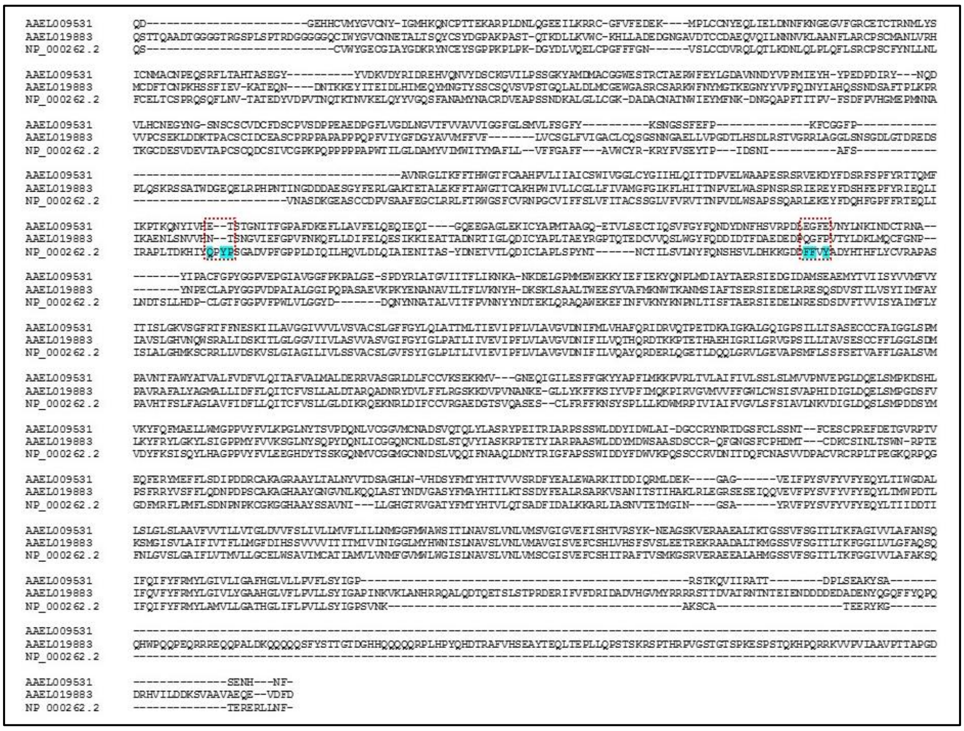

**Fig S4. Sequence alignment of mature *A. aegypti* and *Homo sapiens* NPC1 proteins.** The sequences of mature NPC1 proteins of *A. aegypti* (AAEL009531 and AAEL019883) and Homo sapiens (NP_000262.2) were aligned using Clustal Omega. The six residues of human NPC1 that interact with NPC2 while participating in cholesterol transfer are marked in teal. All of these residues are mutated in *A. aegypti* NPC1 (red dotted boxes).

### Supplementary Tables

#### Table S1. Expression of NPC2 genes in uninfected tissues of *A. aegypti*

##### (A) Expression of NPC2 genes in uninfected midgut

| | Control MG | | |
|---|---|---|---|
| | Day 1* | Day 4# | Day 14§ |
| AAEL001634 | 0.00 | 0.00 | 0.00 |
| AAEL001650 | 365.74 | 1184.16 | 194.51 |
| AAEL001654 | 280.09 | 320.44 | 0.00 |
| AAEL004120 | 336.87 | 195.74 | 659.59 |
| AAEL006854 | 88.65 | 717.35 | 716.79 |
| AAEL007591 | 198676.21 | 252027.37 | 712972.49 |
| AAEL007592 | 26885.00 | 3021.88 | 87141.99 |
| AAEL009553 | 0.00 | 0.00 | 0.00 |
| AAEL009555 | 0.00 | 0.00 | 183.64 |
| AAEL009556 | 162543.71 | 12447.31 | 17460.29 |
| AAEL009760 | 311665.17 | 166901.01 | 664926.92 |
| AAEL009956 | 1887.16 | 161.41 | 844.64 |
| AAEL012064 | 852.68 | 4131.45 | 2686.99 |
| AAEL015136 | 2790862.96 | 1997498.04 | 888536.30 |
| AAEL015137 | 2352.97 | 5564.54 | 9351.43 |
| AAEL015139 | 1152.91 | 0.00 | 1142.74 |
| AAEL015140 | 886.85 | 1776.92 | 190.45 |
| AAEL020314 | 177.40 | 4304.39 | 473.57 |
| AAEL026174 | 49034.14 | 35249.04 | 21863.03 |
| AAEL025109 | 0.00 | 0.00 | 0.00 |

*Each value is the average of 15 individual midguts of Puerto Rico mosquito strain [1].
#Each value is the average of 45 individual midguts of Thailand mosquito strain [2].
§Each value is the average of 10 individual midguts of Mexico mosquito strain [3].

**(B) Expression of NPC2 genes in uninfected salivary glands**

| | Control SG | | |
|---|---|---|---|
| | Day 7* | Day 14*§ | Day 14*§ |
| AAEL001634 | 1.04 | 1.69 | 1.91 |
| AAEL001650 | 0.45 | 0.93 | 1.05 |
| AAEL001654 | 0.00 | 0.00 | 0.00 |
| AAEL004120 | 277.74 | 323.26 | 364.30 |
| AAEL006854 | 7.29 | 14.01 | 15.79 |
| AAEL007591 | 0.45 | 0.55 | 0.62 |
| AAEL007592 | 0.00 | 0.00 | 0.00 |
| AAEL009553 | 0.00 | 0.76 | 0.86 |
| AAEL009555 | 0.00 | 0.38 | 0.43 |
| AAEL009556 | 0.45 | 0.38 | 0.43 |
| AAEL009760 | 1.94 | 5.75 | 6.49 |
| AAEL009956 | 0.00 | 0.00 | 0.00 |
| AAEL012064 | 112.44 | 186.06 | 209.75 |
| AAEL015136 | 0.35 | 0.93 | 1.05 |
| AAEL015137 | 0.00 | 0.55 | 0.62 |
| AAEL015139 | 0.00 | 0.00 | 0.00 |
| AAEL015140 | 0.00 | 0.00 | 0.00 |
| AAEL020314 | 0.00 | 0.00 | 0.00 |
| AAEL026174 | 0.00 | 0.00 | 0.00 |
| AAEL025109 | 0.00 | 0.00 | 0.00 |

*Each value is the average of two independent sets of 20 pairs of salivary glands[4].
§Performed on two independent days [4].

**(C) Expression of NPC2 genes in uninfected antenna, brain, proboscis and whole body**

| | Antenna* | Brain* | Proboscis* | Whole Body# |
|---|---|---|---|---|
| AAEL001634 | 0.00 | 0.00 | 0.00 | 0.00 |
| AAEL001650 | 0.00 | 6724.62 | 0.00 | 0.00 |
| AAEL001654 | 0.00 | 325.62 | 0.00 | 0.00 |
| AAEL004120 | 97325.56 | 255785.10 | 10707.65 | 108777.13 |
| AAEL006854 | 1747848.90 | 17104.74 | 171139.30 | 116420.10 |
| AAEL007591 | 0.00 | 0.00 | 0.00 | 39934.75 |
| AAEL007592 | 825.46 | 0.00 | 0.00 | 27754.65 |
| AAEL009553 | 380.90 | 0.00 | 0.00 | 0.00 |
| AAEL009555 | 0.00 | 0.00 | 0.00 | 0.00 |
| AAEL009556 | 0.00 | 397.33 | 0.00 | 14802.48 |
| AAEL009760 | 419.75 | 0.00 | 0.00 | 1257004.01 |
| AAEL009956 | 0.00 | 0.00 | 0.00 | 0.00 |
| AAEL012064 | 53774.53 | 2060221.71 | 2351248.48 | 416319.74 |
| AAEL015136 | 0.00 | 0.00 | 0.00 | 192459.25 |
| AAEL015137 | 0.00 | 0.00 | 0.00 | 0.00 |
| AAEL015139 | 0.00 | 0.00 | 0.00 | 0.00 |
| AAEL015140 | 0.00 | 0.00 | 0.00 | 0.00 |
| AAEL020314 | 685575.39 | 5704.24 | 74437.94 | 87884.96 |
| AAEL026174 | 0.00 | 0.00 | 0.00 | 1745.58 |
| AAEL025109 | 0.00 | 0.00 | 0.00 | 0.00 |

*Each value is the average of individual sets of 100-220 antenna, 9-18 brain and 275-797 proboscis of Liverpool mosquito strain [5].
#Each value is the average 3 individual whole-body of Liverpool mosquito strain [6].

**Table S2 Highly expressed NPC2 genes in uninfected tissues of *A. aegypti***

**(A) Expression of highly expressed genes in uninfected midgut**

| | Control MG | | | |
|---|---|---|---|---|
| | Day 1* | Day 4# | Day 14§ | |
| AAEL015136 | 1997498.04 | 2790862.96 | 888536.30 | 1 |
| AAEL007591 | 252027.37 | 198676.21 | 712972.49 | 2 |
| AAEL009760 | 166901.01 | 311665.17 | 664926.92 | 3 |
| AAEL026174 | 35249.04 | 49034.14 | 21863.03 | 4 |
| AAEL009556 | 12447.31 | 162543.71 | 17460.29 | 5 |
| AAEL007592 | 3021.88 | 26885.00 | 87141.99 | 6 |
| AAEL015137 | 5564.54 | 2352.97 | 9351.43 | 7 |

*Each value is the average of 15 individual midguts of Puerto Rico mosquito strain [1].
#Each value is the average of 45 individual midguts of Thailand mosquito strain [2].
§Each value is the average of 10 individual midguts of Mexico mosquito strain [3].

**(B) Expression of genes in the uninfected salivary glands**

| | Control SG | | | |
|---|---|---|---|---|
| | Day 7* | Day 14*§ | Day 14*§ | |
| AAEL004120 | 277.74 | 323.26 | 364.30 | 1 |
| AAEL012064 | 112.44 | 186.06 | 209.75 | 2 |
| AAEL006854 | 7.29 | 14.01 | 15.79 | 3 |
| AAEL009760 | 1.94 | 5.75 | 6.49 | 4 |
| AAEL001634 | 1.04 | 1.69 | 1.91 | 5 |
| AAEL001650 | 0.45 | 0.93 | 1.05 | 6 |
| AAEL015136 | 0.35 | 0.93 | 1.05 | 6 |

*Each value is the average of two independent sets of 20 pairs of salivary glands[4].
§Performed on two independent days [4].

**(C) Expression of genes in the uninfected antenna, brain, proboscis and whole body**

| | Antenna* | |
|---|---|---|
| AAEL006854 | 1747848.90 | 1 |
| AAEL020314 | 685575.39 | 2 |
| AAEL004120 | 97325.56 | 3 |
| AAEL012064 | 53774.53 | 4 |

| | Brain* | |
|---|---|---|
| AAEL012064 | 2060221.71 | 1 |
| AAEL004120 | 255785.10 | 2 |
| AAEL006854 | 17104.74 | 3 |
| AAEL001650 | 6724.62 | 4 |
| AAEL020314 | 5704.24 | 5 |

| | Proboscis* | |
|---|---|---|
| AAEL012064 | 2351248.48 | 1 |
| AAEL006854 | 171139.30 | 2 |
| AAEL020314 | 74437.94 | 3 |
| AAEL004120 | 10707.65 | 4 |

| | Whole body# | |
|---|---|---|
| AAEL009760 | 1257004.01 | 1 |
| AAEL012064 | 416319.74 | 2 |
| AAEL015136 | 192459.25 | 3 |
| AAEL006854 | 116420.10 | 4 |
| AAEL004120 | 108777.13 | 5 |
| AAEL020314 | 87884.96 | 6 |

*Each value is the average of individual sets of 100-220 antenna, 9-18 brain and 275-797 proboscis of Liverpool mosquito strain [5].
#Each value is the average 3 individual whole-body of Liverpool mosquito strain [6].

**Table S3 Differential expression in the midgut infected with CHIKV, DENV1 and DENV2**

**(A) Differential expression of genes in the midgut infected with CHIKV**

| | Control MG Day 1 * | CHIKV MG 1-dpi* |
|---|---|---|
| AAEL001634 | 0.00 | 0.00 |
| AAEL001650 | 365.74 | 0.00 |
| AAEL001654 | 280.09 | 0.00 |
| AAEL004120 | 336.87 | 196.51 |
| AAEL006854 | 88.65 | 0.00 |
| AAEL007591 | 198676.21 | 304251.16 |
| AAEL007592 | 26885.00 | 30985.36 |
| AAEL009553 | 0.00 | 0.00 |
| AAEL009555 | 0.00 | 111.09 |
| AAEL009556 | 162543.71 | 114643.93 |
| AAEL009760 | 311665.17 | 427435.53 |
| AAEL009956 | 1887.16 | 2010.28 |
| AAEL012064 | 852.68 | 453.10 |
| AAEL015136 | 2790862.96 | 2695051.43 |
| AAEL015137 | 2352.97 | 3611.56 |
| AAEL015139 | 1152.91 | 1061.27 |
| AAEL015140 | 886.85 | 825.43 |
| AAEL020314 | 177.40 | 353.76 |
| AAEL026174 | 49034.14 | 55193.21 |
| AAEL025109 | 0.00 | 0.00 |

*Each value is the average of 15 individual midguts of Puerto Rico mosquito strain [1].

**(B) Fold change of genes that demonstrate high differential expression in the midgut post-CHIKV infection**

| CHIKV | Down (D) |
|---|---|
| AAEL001650 | D to zero |
| AAEL001654 | D to zero |
| AAEL006854 | D to zero |
| AAEL012064 | 1.9x |
| AAEL004120 | 1.7x |
| | Up (U) |
| AAEL009555 | U from zero |
| AAEL020314 | 2.0x |
| AAEL007591 | 1.5x |
| AAEL015137 | 1.5x |
| AAEL009760 | 1.4x |

D, shut down to Zero expression.
U, initiation of expression from Zero.

**(C) Differential expression of genes in the midgut infected with DENV1**

| | Control MG Day 4 # | DENV1 MG 4-dpi# |
|---|---|---|
| AAEL001634 | 0.00 | 0.00 |
| AAEL001650 | 1184.16 | 0.00 |
| AAEL001654 | 320.44 | 0.00 |
| AAEL004120 | 195.74 | 18.35 |
| AAEL006854 | 717.35 | 50.13 |
| AAEL007591 | 252027.37 | 3303.94 |
| AAEL007592 | 3021.88 | 1751.66 |
| AAEL009553 | 0.00 | 0.00 |
| AAEL009555 | 0.00 | 0.00 |
| AAEL009556 | 12447.31 | 555.57 |
| AAEL009760 | 166901.01 | 2432057.66 |
| AAEL009956 | 161.41 | 0.00 |
| AAEL012064 | 4131.45 | 1020.92 |
| AAEL015136 | 1997498.04 | 36507.28 |
| AAEL015137 | 5564.54 | 73.46 |
| AAEL015139 | 0.00 | 0.00 |
| AAEL015140 | 1776.92 | 72.41 |
| AAEL020314 | 4304.39 | 927.46 |
| AAEL026174 | 35249.04 | 3925.39 |
| AAEL025109 | 0.00 | 0.00 |

#Each value is the average of 45 individual midguts of Thailand mosquito strain [2].

**(D) Fold change of genes that demonstrate high differential expression in the midgut post-DENV1 infection**

| DENV1 | Down (D) |
|---|---|
| AAEL001650 | D to zero |
| AAEL001654 | D to zero |
| AAEL009956 | D to zero |
| AAEL007591 | 76.3x |
| AAEL015137 | 75.7x |
| AAEL015136 | 54.7x |
| AAEL015140 | 24.5x |
| AAEL009556 | 22.4x |
| AAEL006854 | 14.3x |
| AAEL004120 | 10.7x |
| AAEL026174 | 9.0x |
| AAEL020314 | 4.6x |
| AAEL012064 | 4.0x |
| AAEL007592 | 1.7x |
| | Up |
| AAEL009760 | 14.6x |

D, shut down to Zero expression.
U, initiation of expression from Zero.

**(E) Differential expression of genes in the midgut infected with DENV2**

| | Control MG Day 14§ | DENV2 MG 14-dpi§ |
|---|---|---|
| AAEL001634 | 0.00 | 0.00 |
| AAEL001650 | 194.51 | 340.78 |
| AAEL001654 | 0.00 | 0.00 |
| AAEL004120 | 659.59 | 342.40 |
| AAEL006854 | 716.79 | 1171.94 |
| AAEL007591 | 712972.49 | 623372.81 |
| AAEL007592 | 87141.99 | 61405.28 |
| AAEL009553 | 0.00 | 0.00 |
| AAEL009555 | 183.64 | 0.00 |
| AAEL009556 | 17460.29 | 14094.82 |
| AAEL009760 | 664926.92 | 822380.22 |
| AAEL009956 | 844.64 | 696.38 |
| AAEL012064 | 2686.99 | 1113.14 |
| AAEL015136 | 888536.30 | 856842.53 |
| AAEL015137 | 9351.43 | 10471.88 |
| AAEL015139 | 1142.74 | 1048.72 |
| AAEL015140 | 190.45 | 524.32 |
| AAEL020314 | 473.57 | 67.46 |
| AAEL026174 | 21863.03 | 17614.21 |
| AAEL025109 | 0.00 | 0.00 |

§Each value is the average of 10 individual midguts of Mexico mosquito strain [3].

**(F) Fold change of genes that demonstrate high differential expression in the midgut post-DENV2 infection**

| DENV2 | Down (D) |
|---|---|
| AAEL009555 | D to zero |
| AAEL020314 | 7.0x |
| AAEL012064 | 2.4x |
| AAEL004120 | 1.9x |
| AAEL007592 | 1.4x |
| | Up |
| AAEL015140 | 2.8x |
| AAEL001650 | 1.8x |
| AAEL006854 | 1.6x |
| AAEL009760 | 1.2x |

D, shut down to Zero expression.
U, initiation of expression from Zero.

**Table S4 Differential expression in salivary glands infected with CHIKV, DENV2 and ZIKV**

**(A) Differential expression of genes in the salivary gland infected with CHIKV**

| | Control SG Day 7* | CHIKV SG 7-dpi* |
|---|---|---|
| AAEL001634 | 1.04 | 0.00 |
| AAEL001650 | 0.45 | 5.54 |
| AAEL001654 | 0.00 | 0.00 |
| AAEL004120 | 277.74 | 272.52 |
| AAEL006854 | 7.29 | 6.69 |
| AAEL007591 | 0.45 | 0.00 |
| AAEL007592 | 0.00 | 0.00 |
| AAEL009553 | 0.00 | 0.00 |
| AAEL009555 | 0.00 | 0.00 |
| AAEL009556 | 0.45 | 0.00 |
| AAEL009760 | 1.94 | 1.41 |
| AAEL009956 | 0.00 | 0.00 |
| AAEL012064 | 112.44 | 84.78 |
| AAEL015136 | 0.35 | 0.00 |
| AAEL015137 | 0.00 | 0.00 |
| AAEL015139 | 0.00 | 0.00 |
| AAEL015140 | 0.00 | 0.00 |

* Each value is the average of two independent sets of 20 pairs of salivary glands [4].
§Performed on two independent days [4].

**(B) Fold change of genes that demonstrate high differential expression in the salivary gland post-CHIKV infection**

| CHIKV | Down (D) |
|---|---|
| AAEL001634 | D to zero |
| AAEL007591 | D to zero |
| AAEL009760 | 1.4x |
| AAEL012064 | 1.3x |
| AAEL006854 | 1.1x |
| AAEL004120 | 1.0x |
| AAEL001650 | 12.3x |

D, shut down to Zero expression.
U, initiation of expression from Zero.

**(C) Differential expression of genes in the salivary gland infected with DENV2**

| | Control SG Day 14* | DENV2 SG 14-dpi * |
|---|---|---|
| AAEL001634 | 1.69 | 0.48 |
| AAEL001650 | 0.93 | 4.72 |
| AAEL001654 | 0.00 | 0.00 |
| AAEL004120 | 323.26 | 195.34 |
| AAEL006854 | 14.01 | 5.09 |
| AAEL007591 | 0.55 | 0.00 |
| AAEL007592 | 0.00 | 0.00 |
| AAEL009553 | 0.76 | 0.00 |
| AAEL009555 | 0.38 | 0.48 |
| AAEL009556 | 0.38 | 2.43 |
| AAEL009760 | 5.75 | 5.69 |
| AAEL009956 | 0.00 | 0.00 |
| AAEL012064 | 186.06 | 126.29 |
| AAEL015136 | 0.93 | 0.00 |
| AAEL015137 | 0.55 | 0.00 |
| AAEL015139 | 0.00 | 0.00 |
| AAEL015140 | 0.00 | 0.00 |

* Each value is the average of two independent sets of 20 pairs of salivary glands [4].
§Performed on two independent days [4].

**(D) Fold change of genes that demonstrate high differential expression in the salivary gland post-DENV2 infection**

| DENV2 | Down (D) |
|---|---|
| AAEL007591 | D to zero |
| AAEL009553 | D to zero |
| AAEL015136 | D to zero |
| AAEL015137 | D to zero |
| AAEL001634 | 3.5x |
| AAEL006854 | 2.8x |
| AAEL004120 | 1.7x |
| AAEL012064 | 1.5x |
| | Up |
| AAEL009556 | 6.4x |
| AAEL001650 | 5.1x |
| AAEL009555 | 1.3x |

D, shut down to Zero expression.
U, initiation of expression from Zero.

**(E) Differential expression of genes in the salivary gland infected with ZIKV**

| | Control SG Day 14* | ZIKV SG 14-dpi* |
|---|---|---|
| AAEL001634 | 1.91 | 1.40 |
| AAEL001650 | 1.05 | 1.40 |
| AAEL001654 | 0.00 | 0.00 |
| AAEL004120 | 364.30 | 394.85 |
| AAEL006854 | 15.79 | 24.11 |
| AAEL007591 | 0.62 | 1.61 |
| AAEL007592 | 0.00 | 4.29 |
| AAEL009553 | 0.86 | 0.00 |
| AAEL009555 | 0.43 | 0.00 |
| AAEL009556 | 0.43 | 0.00 |
| AAEL009760 | 6.49 | 2.47 |
| AAEL009956 | 0.00 | 0.00 |
| AAEL012064 | 209.75 | 129.77 |
| AAEL015136 | 1.05 | 1.07 |
| AAEL015137 | 0.62 | 0.00 |
| AAEL015139 | 0.00 | 0.00 |
| AAEL015140 | 0.00 | 0.00 |

* Each value is the average of two independent sets of 20 pairs of salivary glands [4].
§Performed on two independent days [4].

**(F) Fold change of genes that demonstrate high differential expression in the salivary gland post-ZIKV infection**

| ZIKV | Down (D) |
|---|---|
| AAEL009553 | D to zero |
| AAEL009555 | D to zero |
| AAEL009556 | D to zero |
| AAEL015137 | D to zero |
| AAEL009760 | 2.6x |
| AAEL012064 | 1.6x |
| AAEL001634 | 1.4x |
| | Up (U) |
| AAEL007592 | U from zero |
| AAEL007591 | 2.6x |
| AAEL006854 | 1.5x |
| AAEL001650 | 1.3x |

D, shut down to Zero expression.
U, initiation of expression from Zero.

### Table S5 Differential expression of TFs of group A genes in the uninfected and infected (CHIKV, DENV1 and DENV2) midgut

#### (A) Differential expression of TFs of group A genes expressed exclusively in the midgut of uninfected and infected (CHIKV, DENV1 and DENV2) mosquitos

| TFs common to AAEL006854,AAEL020314 | | | | | | | | | | | | |
|---|---|---|---|---|---|---|---|---|---|---|---|---|
| | Drosophila homolog | | Ae. aegypti Gen | Control (1 day) | CHIKV (1 day) | % | Control (4 days) | DENV1 (4 days) | % | Control (14 days | DENV2 (14days) | % |
| Sp1 | Sp1 | A8JUN6_DROME | AAEL003875 | 8.82 | 8.25 | -6.46 | 2.68 | 2.85 | 6.34 | 9.81 | 7.24 | -26.20 |
| Oct-1 | Nub | PDM1_DROME | AAEL017445 | 27.24 | 25.08 | -7.93 | 2.07 | 1.97 | -4.83 | 25.22 | 15.41 | -38.91 |
| HNF-1 | HNF-1 | HNF1_DROME | AAEL011323 | 62.50 | 59.91 | -4.14 | 62.04 | 69.13 | 11.43 | 49.43 | 62.65 | 26.76 |
| C/EBPbeta | Slbo | CEBP_DROME | AAEL002853 | 46.54 | 36.03 | -22.58 | 35.34 | 29.64 | -16.13 | 54.12 | 39.25 | -27.47 |
| GLO | GLO | Q8IG99_DROME | AAEL005947 | 66.09 | 57.49 | -13.01 | 18.92 | 25.87 | 36.73 | 39.78 | 41.00 | 3.07 |
| CRE-BP1 | CREB | CREBA_DROME | AAEL000402 | 4.01 | 3.32 | -17.21 | 1.85 | 2.51 | 35.68 | 2.99 | 1.23 | -58.84 |
| AP1 | AP1 | AP1_DROME | AAEL005364 | 39.1 | 35.08 | -10.28 | 19.48 | 26.14 | 34.19 | 24.24 | 19.35 | -20.17 |
| c-Jun | Jun | JUN_DROME | AAEL003505 | 92.07 | 80.36 | -12.72 | 19.77 | 30.15 | 52.50 | 69.20 | 41.66 | -39.80 |
| c-Fos | Fos | FOS_DROME | AAEL008953 | 51.48 | 49.01 | -4.80 | 4.35 | 7.61 | 74.94 | 118.65 | 68.24 | -42.48 |
| Odd | Odd | ODD_DROME | AAEL007450 | 14.36 | 13.47 | -6.20 | 3.28 | 9.14 | 178.66 | 10.97 | 7.59 | -30.78 |
| Erg | Pnt | PNT_DROME | AAEL003845 | 11.54 | 10.91 | -5.46 | 0.62 | 0.73 | 17.74 | 16.69 | 10.83 | -35.10 |
| TFs unique to AAEL020314 | | | | | | | | | | | | |
| | Drosophila homolog | | Ae. aegypti Gen | Control (1 day) | CHIKV (1 day) | % | Control (4 days) | DENV1 (4 days) | % | Control (14 days | DENV2 (14days) | % |
| Hb | Hb | HUNB_DROME | AAEL000894 | 0 | 0 | | 0.01 | 0.00 | | 0.0 | 0.0 | |
| TFs unique to AAEL006854 | | | | | | | | | | | | |
| | Drosophila homolog | | Ae. aegypti Gen | Control (1 day) | CHIKV (1 day) | % | Control (4 days) | DENV1 (4 days) | % | Control (14 days | DENV2 (14days) | % |
| Nf-kB | NFkB | NFKB1_DROME | AAEL007624 | 21.33 | 20.75 | -2.72 | 6.20 | 10.62 | 71.29 | 26.85 | 19.22 | -28.42 |
| TBP | TBP | TBP_DROME | AAEL012330 | 8.12 | 7.59 | -6.53 | 7.74 | 9.44 | 21.96 | 5.83 | 7.81 | 33.95 |

[1-3]

TFs are divided into three sections i.e., common to both genes, unique to AAEL006854 and unique to AAEL020314.

**(B) Differential expression of TFs that show >20% change in the midgut post-CHIKV, -DENV1 and -DENV2.**

| | | | | |
|---|---|---|---|---|
| | | | Down | Up |
| Control | CHIKV MG | % | AAEL006854 | AAEL020314 |
| 46.54 | 36.03 | -22.58 | C/EBPbeta | C/EBPbeta |
| | | | Down | Down |
| Control | DENV1 MG | % | AAEL006854 | AAEL020314 |
| 18.92 | 25.87 | 36.73 | GLO | GLO |
| 1.85 | 2.51 | 35.68 | CRE-BP1 | CRE-BP1 |
| 19.48 | 26.14 | 34.19 | AP1 | AP1 |
| 19.77 | 30.15 | 52.50 | c-Jun | c-Jun |
| 4.35 | 7.61 | 74.94 | c-Fos | c-Fos |
| 3.28 | 9.14 | 178.66 | Odd | Odd |
| 6.20 | 10.62 | 71.29 | NF-Kb | |
| 7.74 | 9.44 | 21.96 | TBP | |
| | | | Up | Down |
| Control | DENV2 MG | % | AAEL006854 | AAEL020314 |
| 49.43 | 62.65 | 26.76 | HNF-1 | HNF-1 |
| 5.83 | 7.81 | 33.95 | TBP | |
| 9.81 | 7.24 | -26.20 | Sp1 | Sp1 |
| 25.22 | 15.41 | -38.91 | Oct-1 | Oct-1 |
| 54.12 | 39.25 | -27.47 | C/EBPbeta | C/EBPbeta |
| 2.99 | 1.23 | -58.84 | CRE-BP1 | CRE-BP1 |
| 24.24 | 19.35 | -20.17 | AP1 | AP1 |
| 69.20 | 41.66 | -39.80 | c-Jun | c-Jun |
| 118.65 | 68.24 | -42.48 | c-Fos | c-Fos |
| 10.97 | 7.59 | -30.78 | Odd | Odd |
| 16.69 | 10.83 | -35.10 | Erg | Erg |
| 26.85 | 19.22 | -28.42 | NF-kB | |

The TFs that show an upregulation and downregulation are indicated with yellow and purple, respectively.

#### Table S6 Differential expression of TFs of group A genes in the uninfected and infected (CHIKV, DENV2 and ZIKV) salivary glands

#### (A) Differential expression of TFs of group A genes expressed exclusively in the salivary glands of uninfected and infected (CHIKV, DENV2 and ZIKV) mosquitos

| TFs common to AAEL006854,AAEL020314 | | | | | | | | | | | |
|---|---|---|---|---|---|---|---|---|---|---|---|
| | Drosophila homolog | | Ae. aegypti Gen | Control (7 days) | CHIKV (7 day) | % | Control (14 days | DENV2 (14 days | % | ZIKV (14 days) | % |
| Sp1 | Sp1 | A8JUN6_DROME | AAEL003875 | 205.08 | 270.14 | 31.72 | 280.37 | 266.65 | -3.67 | 332.58 | 18.62 |
| HNF-1 | HNF-1 | HNF1_DROME | AAEL011323 | 375.31 | 424.07 | 12.99 | 433.75 | 454.67 | 8.86 | 591.66 | 36.41 |
| C/EBPbeta | Slbo | CEBP_DROME | AAEL002853 | 750.94 | 524.27 | -30.18 | 706.59 | 692.82 | -4.25 | 637.88 | -9.72 |
| GLO | GLO | Q8IG99_DROME | AAEL005947 | 339.97 | 350.87 | 3.21 | 406.43 | 434.89 | -3.71 | 482.19 | 18.64 |
| CRE-BP1 | CREB | CREBA_DROME | AAEL000402 | 123.87 | 125.23 | 1.09 | 214.89 | 155.30 | -14.15 | 221.07 | 2.88 |
| c-Jun | Jun | JUN_DROME | AAEL003505 | 478.61 | 1196.77 | 150.05 | 645.06 | 854.29 | -5.43 | 690.33 | 7.02 |
| c-Fos | Fos | FOS_DROME | AAEL008953 | 748.49 | 1099.60 | 46.91 | 1107.61 | 1363.26 | -5.72 | 1322.66 | 19.42 |
| Erg | Pnt | PNT_DROME | AAEL003845 | 27.99 | 41.19 | 47.15 | 54.48 | 49.24 | -24.90 | 63.61 | 16.76 |
| TFs unique to AAEL020314 | | | | | | | | | | | |
| | Drosophila homolog | | Ae. aegypti Gen | Control (7 days) | CHIKV (7 day) | % | Control (14 days | DENV2 (14 days | % | ZIKV (14 days) | % |
| Hb | Hb | HUNB_DROME | AAEL000894 | 0.00 | 0.00 | 0.00 | 0.00 | 0.00 | 0.00 | 0.00 | 0.00 |
| TFs unique to AAEL006854 | | | | | | | | | | | |
| | Drosophila homolog | | Ae. aegypti Gen | Control (7 days) | CHIKV (7 day) | % | Control (14 days | DENV2 (14 days | % | ZIKV (14 days) | % |
| Nf-kB | NFkB | NFKB1_DROME | AAEL007624 | 339.55 | 414.48 | 22.07 | 509.17 | 476.07 | -6.50 | 599.33 | 17.71 |
| TBP | TBP | TBP_DROME | AAEL012330 | 61.71 | 54.77 | -11.25 | 69.54 | 85.09 | 22.36 | 98.81 | 42.08 |

[4]

TFs are divided into three sections i.e., common to both genes, unique to AAEL006854 and unique to AAEL020314.

**(B) Differential expression of TFs that show >20% change in the salivary glands post-CHIKV, -DENV2 and -ZIKV**

| | | | No change | No exp |
|---|---|---|---|---|
| Control | CHIKV SG | % | AAEL006854 | AAEL020314 |
| 205.08 | 270.14 | 31.72 | sp1 | sp1 |
| 478.61 | 1196.77 | 150.05 | Jun | Jun |
| 748.49 | 1099.60 | 46.91 | Fos | Fos |
| 27.99 | 41.19 | 47.15 | ERG | ERG |
| 339.55 | 414.48 | 22.07 | nf-kB | |
| 750.94 | 524.27 | -30.18 | CEBP | CEBP |
| | | | Down | No exp |
| Control | DENV2 SG | % | AAEL006854 | AAEL020314 |
| 69.54 | 85.09 | 22.36 | TBP | |
| 54.48 | 49.24 | -24.90 | ERG | ERG |
| | | | Up | No exp |
| Control | ZIKV SG | % | AAEL006854 | AAEL020314 |
| 433.75 | 591.66 | 36.41 | HNF-1 | HNF-1 |
| 69.54 | 98.81 | 42.08 | TBP | |

The TFs that show an upregulation and downregulation are indicated with yellow and purple, respectively.

### Table S7 Differential expression of TFs of group B genes in the uninfected and infected (CHIKV, DENV1 and DENV2) midgut

#### (A) Differential expression of TFs of group B genes expressed exclusively in the midgut of uninfected and infected (CHIKV, DENV1 and DENV2) mosquitos

| | | | | | | | | | | | | |
|---|---|---|---|---|---|---|---|---|---|---|---|---|
| TFs common to AAEL009553, AAEL009555, AAEL009556 | | | | | | | | | | | | |
| | Drosophila homolog | | Ae. aegypti Gene | Control (1 day) | CHIKV (1 day) | % | Control (4 days) | DENV1 (4 days) | % | Control (14 days | DENV2 (14days) | % |
| Sp1 | Sp1 | A8JUN6_DROME | AAEL003875 | 8.82 | 8.25 | -6.46 | 2.68 | 2.85 | 6.34 | 9.81 | 7.24 | -26.20 |
| Oct-1 | Nub | PDM1_DROME | AAEL017445 | 27.24 | 25.08 | -7.93 | 2.07 | 1.97 | -4.83 | 25.22 | 15.41 | -38.91 |
| HNF-1 | HNF1 | HNF1_DROME | AAEL011323 | 62.50 | 59.91 | -4.14 | 62.04 | 69.13 | 11.43 | 49.43 | 62.65 | 26.76 |
| YY1 | Pho | PHO_DROME | AAEL008005 | 13.27 | 12.10 | -8.82 | 5.26 | 8.86 | 68.44 | 8.27 | 5.80 | -29.90 |
| MCM1 | MCM1 | MCM1_DROME | AAEL007007 | 2.10 | 2.03 | -3.33 | 0.50 | 1.31 | 162.00 | 2.87 | 1.24 | -56.76 |
| TBP | TBP | TBP_DROME | AAEL012330 | 8.12 | 7.59 | -6.53 | 7.74 | 9.44 | 21.96 | 5.83 | 7.81 | 33.95 |
| TFs common to AAEL009553, AAEL009555 | | | | | | | | | | | | |
| | Drosophila homolog | | Ae. aegypti Gene | Control (1 day) | CHIKV (1 day) | | Control (4 days) | DENV1 (4 days) | | Control (14 days | DENV2 (14days) | |
| RAP1 | RAP1 | RAP1_DROME | AAEL009377 | 48.88 | 44.38 | -9.21 | 20.90 | 31.28 | 49.67 | 29.08 | 22.04 | -24.21 |
| NF-kB | NFkB | NFKB1_DROME | AAEL007624 | 21.33 | 20.75 | -2.72 | 6.20 | 10.62 | 71.29 | 26.85 | 19.22 | -28.42 |
| TFs common to AAEL9553, AAEL009556 | | | | | | | | | | | | |
| | Drosophila homolog | | Ae. aegypti Gene | Control (1 day) | CHIKV (1 day) | | Control (4 days) | DENV1 (4 days) | | Control (14 days | DENV2 (14days) | |
| C/EBPbeta | Slbo | CEBP_DROME | AAEL002853 | 46.54 | 36.03 | -22.58 | 35.34 | 29.64 | -16.13 | 54.12 | 39.25 | -27.47 |
| TFs common to AAEL009555, AAEL009556 | | | | | | | | | | | | |
| | Drosophila homolog | | Ae. aegypti Gene | Control (1 day) | CHIKV (1 day) | | Control (4 days) | DENV1 (4 days) | | Control (14 days | DENV2 (14days) | |
| Ftz | Ftz | FTZ_DROME | AAEL009947 | 1.71 | 1.68 | -1.75 | 0.82 | 0.90 | 9.76 | 0.96 | 0.61 | -36.26 |
| Erg | Pnt | PNT_DROME | AAEL003845 | 11.54 | 10.91 | -5.46 | 0.62 | 0.73 | 17.74 | 16.69 | 10.83 | -35.10 |
| TFs unique to AAEL009556 | | | | | | | | | | | | |
| | Drosophila homolog | | Ae. aegypti Gene | Control (1 day) | CHIKV (1 day) | | Control (4 days) | DENV1 (4 days) | | Control (14 days | DENV2 (14days) | |
| USF | USF | Q9W4J8_DROME | AAEL006314 | 3.03 | 2.33 | -23.10 | 4.04 | 6.64 | 64.36 | 3.47 | 3.54 | 2.05 |
| NF-Atc | NFAT | E1JJP5_DROME | AAEL011359 | 2.28 | 1.99 | -12.72 | 0.68 | 0.50 | -26.47 | 0.76 | 0.49 | -35.12 |
| RXR-beta/alpha | USP | USP_DROME | AAEL000395 | 17.25 | 21.76 | 26.14 | 1.52 | 2.07 | 36.18 | 3.93 | 3.38 | -13.83 |
| TFs unique to AAEL009553 | | | | | | | | | | | | |
| | Drosophila homolog | | Ae. aegypti Gene | Control (1 day) | CHIKV (1 day) | | Control (4 days) | DENV1 (4 days) | | Control (14 days | DENV2 (14days) | |
| SRF | Bs | SRF_DROME | AAEL008721 | 2.36 | 2.03 | -13.98 | 2.02 | 2.29 | 13.37 | 1.22 | 0.77 | -36.58 |
| HSE-Bind | HSF | HSF_DROME | AAEL010319 | 23.75 | 23.11 | -2.69 | 11.51 | 13.70 | 19.03 | 18.87 | 9.56 | -49.31 |
| Pit-1a | Pita | Q95RQ8_DROME | AAEL003529 | 2.95 | 2.77 | -6.10 | 2.23 | 2.96 | 32.74 | 1.83 | 2.40 | 30.56 |
| GLO | GLO | Q8IG99_DROME | AAEL005947 | 66.09 | 57.49 | -13.01 | 18.92 | 25.87 | 36.73 | 39.78 | 41.00 | 3.07 |
| TFs unique to AAEL009555 | | | | | | | | | | | | |
| | Drosophila homolog | | Ae. aegypti Gene | Control (1 day) | CHIKV (1 day) | | Control (4 days) | DENV1 (4 days) | | Control (14 days | DENV2 (14days) | |
| CRE-BP1 | CREB | CREBA_DROME | AAEL000402 | 4.01 | 3.32 | -17.21 | 1.85 | 2.51 | 35.68 | 2.99 | 1.23 | -58.84 |
| SGF | SGF | FKH_DROME | AAEL003163 | 3.81 | 3.40 | -10.76 | 0.18 | 0.30 | 66.67 | 3.81 | 1.92 | -49.55 |
| GABP | ELG | ELG_DROME | AAEL009584 | 8.76 | 7.57 | -13.58 | 3.37 | 4.44 | 31.75 | 3.97 | 4.14 | 4.37 |

[1-3]

TFs are divided into seven sections. The first section is common to all genes, the next three are common to any two genes, and the last three sections are unique to each gene.

**(B) Differential expression of TFs that show >20% change in the midgut post-CHIKV, -DENV1 and -DENV2**

| | | | No expression | Up | Down |
|---|---|---|---|---|---|
| **Control** | **CHIKV MG** | **%** | **AAEL009553** | **AAEL009555** | **AAEL009556** |
| 17.25 | 21.76 | 26.14 | | | RXR-beta/alpha |
| 46.54 | 36.03 | -22.58 | C/EBPbeta | | C/EBPbeta |
| 3.03 | 2.33 | -23.10 | | | USF |
| | | | No expression | No expression | Down |
| **Control** | **DENV1 MG** | **%** | **AAEL009553** | **AAEL009555** | **AAEL009556** |
| 5.26 | 8.86 | 68.44 | YY1 | YY1 | YY1 |
| 0.50 | 1.31 | 162.00 | MCM1 | MCM1 | MCM1 |
| 7.74 | 9.44 | 21.96 | TBP | TBP | TBP |
| 20.90 | 31.28 | 49.67 | RAP1 | RAP1 | |
| 6.20 | 10.62 | 71.29 | Nf-kB | Nf-kB | |
| 2.23 | 2.96 | 32.74 | Pit-1a | | |
| 18.92 | 25.87 | 36.73 | GLO | | |
| 1.85 | 2.51 | 35.68 | | CRE-BP1 | |
| 0.18 | 0.30 | 66.67 | | SGF | |
| 3.37 | 4.44 | 31.75 | | GABP | |
| 4.04 | 6.64 | 64.36 | | | USF |
| 1.52 | 2.07 | 36.18 | | | RXR-beta/alpha |
| 0.68 | 0.50 | -26.47 | | | NF-Atc |
| | | | No expression | Down | No change |
| **Control** | **DENV2 MG** | **%** | **AAEL009553** | **AAEL009555** | **AAEL009556** |
| 49.43 | 62.65 | 26.76 | HNF-1 | HNF-1 | HNF-1 |
| 5.83 | 7.81 | 33.95 | TBP | TBP | TBP |
| 1.83 | 2.40 | 30.56 | Pit-1a | | |
| 9.81 | 7.24 | -26.20 | Sp1 | Sp1 | Sp1 |
| 25.22 | 15.41 | -38.91 | Oct-1 | Oct-1 | Oct-1 |
| 8.27 | 5.80 | -29.90 | YY1 | YY1 | YY1 |
| 2.87 | 1.24 | -56.76 | MCM1 | MCM1 | MCM1 |
| 29.08 | 22.04 | -24.21 | RAP1 | RAP1 | |
| 26.85 | 19.22 | -28.42 | Nf-kB | Nf-kB | |
| 0.96 | 0.61 | -36.26 | | Ftz | Ftz |
| 16.69 | 10.83 | -35.10 | | Erg | Erg |
| 54.12 | 39.25 | -27.47 | C/EBPbeta | | C/EBPbeta |
| 1.22 | 0.77 | -36.58 | SRF | | |
| 18.87 | 9.56 | -49.31 | HSE-Bind | | |
| 2.99 | 1.23 | -58.84 | | CRE-BP1 | |
| 3.81 | 1.92 | -49.55 | | SGF | |
| 0.76 | 0.49 | -35.12 | | | NF-Atc |

The TFs that show an upregulation and downregulation are indicated with yellow and purple, respectively.

#### Table S8 Differential expression of TFs of group B genes in the uninfected and infected (CHIKV, DENV2 and ZIKV) salivary glands

#### (A) Differential expression of TFs of group B genes expressed exclusively in the salivary glands of uninfected and infected (CHIKV, DENV1 and DENV2) mosquitos

| | | | | | | | | | | | | |
|---|---|---|---|---|---|---|---|---|---|---|---|---|
| TFs common to AAEL009553, AAEL009555, AAEL009556 | | | | | | | | | | | | |
| | Drosophila homolog | | Ae. aegypti Gen | Control (7 day) | CHIKV (7 day) | % | Control ( 14 day | DENV2 (14 day) | % | Control ( 14 day | ZIKV (14 day) | % |
| Sp1 | Sp1 | A8JUN6_DROME | AAEL003875 | 205.08 | 270.14 | 31.72 | 280.37 | 266.65 | -4.89 | 280.37 | 332.58 | 18.62 |
| HNF-1 | HNF1 | HNF1_DROME | AAEL011323 | 375.31 | 424.07 | 12.99 | 433.75 | 454.67 | 4.82 | 433.75 | 591.66 | 36.41 |
| YY1 | Pho | PHO_DROME | AAEL008005 | 28.97 | 41.96 | 44.83 | 63.53 | 54.52 | -14.17 | 63.53 | 64.80 | 2.01 |
| MCM1 | MCM1 | MCM1_DROME | AAEL007007 | 23.31 | 78.81 | 238.07 | 35.48 | 75.38 | 112.47 | 35.48 | 52.12 | 46.91 |
| TBP | TBP | TBP_DROME | AAEL012330 | 61.71 | 54.77 | -11.25 | 69.54 | 85.09 | 22.36 | 69.54 | 98.81 | 42.08 |
| TFs common to AAEL009553, AAEL009555 | | | | | | | | | | | | |
| | Drosophila homolog | | Ae. aegypti Gen | Control (7 day) | CHIKV (7 day) | | Control ( 14 day | DENV2 (14 day) | | Control ( 14 day | ZIKV (14 day) | |
| RAP1 | RAP1 | RAP1_DROME | AAEL009377 | 247.26 | 252.88 | 2.27 | 345.48 | 321.01 | -7.08 | 345.48 | 341.81 | -1.06 |
| NF-kB | NFkB | NFKB1_DROME | AAEL007624 | 339.55 | 414.48 | 22.07 | 509.17 | 476.07 | -6.50 | 509.17 | 599.33 | 17.71 |
| TFs common to AAEL9553, AAEL009556 | | | | | | | | | | | | |
| | Drosophila homolog | | Ae. aegypti Gen | Control (7 day) | CHIKV (7 day) | | Control ( 14 day | DENV2 (14 day) | | Control ( 14 day | ZIKV (14 day) | |
| C/EBPbeta | Slbo | CEBP_DROME | AAEL002853 | 750.94 | 524.27 | -30.18 | 706.59 | 692.82 | -1.95 | 706.59 | 637.88 | -9.72 |
| TFs common to AAEL009555, AAEL009556 | | | | | | | | | | | | |
| | Drosophila homolog | | Ae. aegypti Gen | Control (7 day) | CHIKV (7 day) | | Control ( 14 day | DENV2 (14 day) | | Control ( 14 day | ZIKV (14 day) | |
| Ftz | Ftz | FTZ_DROME | AAEL009947 | 14.14 | 9.46 | -33.08 | 15.33 | 6.06 | -60.45 | 15.33 | 13.35 | -12.90 |
| COUP | Svp | A0B4KFQ2_DROI | AAEL002765 | 16.44 | 19.31 | 17.41 | 24.35 | 6.79 | -72.11 | 24.35 | 24.65 | 1.21 |
| Erg | Pnt | PNT_DROME | AAEL003845 | 27.99 | 41.19 | 47.15 | 54.48 | 49.24 | -9.63 | 54.48 | 63.61 | 16.76 |
| TFs unique to AAEL009556 | | | | | | | | | | | | |
| | Drosophila homolog | | Ae. aegypti Gen | Control (7 day) | CHIKV (7 day) | | Control ( 14 day | DENV2 (14 day) | | Control ( 14 day | ZIKV (14 day) | |
| USF | USF | Q9W4J8_DROME | AAEL006314 | 133.21 | 102.57 | -23.00 | 153.79 | 170.57 | 10.92 | 153.79 | 244.82 | 59.20 |
| NF-Atc | NFAT | E1JJP5_DROME | AAEL011359 | 101.62 | 78.49 | -22.76 | 160.51 | 133.18 | -17.02 | 160.51 | 180.36 | 12.37 |
| RXR-beta/alpha | USP | USP_DROME | AAEL000395 | 156.93 | 120.88 | -22.97 | 207.82 | 226.48 | 8.98 | 207.82 | 236.80 | 13.94 |
| ER | ERR | Q8WS79_DROME | AAEL013546 | 304.50 | 298.58 | -1.94 | 449.70 | 329.51 | -26.73 | 449.70 | 434.10 | -3.47 |
| TFs unique to AAEL009553 | | | | | | | | | | | | |
| | Drosophila homolog | | Ae. aegypti Gen | Control (7 day) | CHIKV (7 day) | | Control ( 14 day | DENV2 (14 day) | | Control ( 14 day | ZIKV (14 day) | |
| HSE-Bind | HSF | HSF_DROME | AAEL010319 | 302.36 | 276.07 | -8.70 | 347.17 | 346.73 | -0.13 | 347.17 | 390.37 | 12.44 |
| Pit-1a | Pita | Q95RQ8_DROME | AAEL003529 | 29.70 | 25.96 | -12.59 | 49.37 | 52.61 | 6.55 | 49.37 | 60.08 | 21.68 |
| GLO | GLO | Q8IG99_DROME | AAEL005947 | 339.97 | 350.87 | 3.21 | 406.43 | 434.89 | 7.00 | 406.43 | 482.19 | 18.64 |
| TFs unique to AAEL009555 | | | | | | | | | | | | |
| | Drosophila homolog | | Ae. aegypti Gen | Control (7 day) | CHIKV (7 day) | | Control ( 14 day | DENV2 (14 day) | | Control ( 14 day | ZIKV (14 day) | |
| TAF-1 | TAF1 | TAF1_DROME | AAEL014023 | 68.97 | 59.39 | -13.88 | 64.15 | 73.42 | 14.46 | 64.15 | 81.82 | 27.54 |
| CRE-BP1 | CREB | CREBA_DROME | AAEL000402 | 123.87 | 125.23 | 1.09 | 214.89 | 155.30 | -27.73 | 214.89 | 221.07 | 2.88 |
| SGF | SGF | FKH_DROME | AAEL003163 | 186.73 | 225.56 | 20.79 | 293.60 | 415.14 | 41.40 | 293.60 | 338.98 | 15.46 |
| GABP | ELG | ELG_DROME | AAEL009584 | 54.89 | 33.76 | -38.50 | 78.86 | 73.68 | -6.57 | 78.86 | 73.22 | -7.15 |

[4]

TFs are divided into seven sections. The first section is common to all genes, the next three are common to any two genes, and the last three sections are unique to each gene.

**(B) Differential expression of TFs that show >20% change in the salivary glands post- CHIKV, -DENV2 and -ZIKV.**

| | | | No expression | No expression | Down |
|---|---|---|---|---|---|
| Control | CHIKV SG | % | AAEL009553 | AAEL009555 | AAEL009556 |
| 205.08 | 270.14 | 31.72 | Sp1 | Sp1 | Sp1 |
| 28.97 | 41.96 | 44.83 | YY1 | YY1 | YY1 |
| 23.31 | 78.81 | 238.07 | MCM1 | MCM1 | MCM1 |
| 339.55 | 414.48 | 22.07 | Nf-kB | Nf-kB | |
| 27.99 | 41.19 | 47.15 | | ERG | ERG |
| 186.73 | 225.56 | 20.79 | | SGF | |
| 750.94 | 524.27 | -30.18 | CEBP | | CEBP |
| 14.14 | 9.46 | -33.08 | | Ftz | Ftz |
| 54.89 | 33.76 | -38.50 | | GABP | |
| 133.21 | 102.57 | -23.00 | | | USF |
| 101.62 | 78.49 | -22.76 | | | NF-Atc |
| 156.93 | 120.88 | -22.97 | | | RXR-beta/alpha |
| | | | Down | Up | Up |
| Control | DENV2 SG | % | AAEL009553 | AAEL009555 | AAEL009556 |
| 35.48 | 75.38 | 112.47 | MCM1 | MCM1 | MCM1 |
| 69.54 | 85.09 | 22.36 | TBP | TBP | TBP |
| 293.60 | 415.14 | 41.40 | | SGF | |
| 15.33 | 6.06 | -60.45 | | Ftz | Ftz |
| 24.35 | 6.79 | -72.11 | | COUP | COUP |
| 214.89 | 155.30 | -27.73 | | CRE-BP1 | |
| 449.70 | 329.51 | -26.73 | | | ER |
| | | | Down | Down | Down |
| Control | ZIKV SG | % | AAEL009553 | AAEL009555 | AAEL009556 |
| 433.75 | 591.66 | 36.41 | HNF-1 | HNF-1 | HNF-1 |
| 35.48 | 52.12 | 46.91 | MCM1 | MCM1 | MCM1 |
| 69.54 | 98.81 | 42.08 | TBP | TBP | TBP |
| 49.37 | 60.08 | 21.68 | Pita | | |
| 64.15 | 81.82 | 27.54 | | TAF1 | |
| 153.79 | 244.82 | 59.20 | | | USF |

The TFs that show an upregulation and downregulation are indicated with yellow and purple, respectively.
